# Ternary complex formation of VCP, VCPIP1 and p47 facilitates Golgi reassembly

**DOI:** 10.1101/2024.10.01.616120

**Authors:** Binita Shah, Moritz Hunkeler, Ariana Bratt, Hong Yue, Isabella Jaen Maisonet, Eric S. Fischer, Sara J. Buhrlage

## Abstract

VCP/p97 regulates a wide range of cellular processes, including post-mitotic Golgi reassembly. In this context, VCP is assisted by p47, an adapter protein, and VCPIP1, a deubiquitinase (DUB). However, how they organize into a functionally competent complex and promote reassembly remains unknown. Here, we use cryo-EM to characterize VCP-VCPIP1 and VCP-VCPIP1-p47 complexes. We show that VCPIP1 engages VCP through two interfaces: one involving the N-domain of VCP and the UBX domain of VCPIP1, and the other involving the VCP D2 domains and a region of VCPIP1 we refer to as VCPID. The p47 UBX domain competitively binds to the VCP N-domain, while not affecting VCPID binding. We show that VCPID is critical for VCP-mediated enhancement of DUB activity and Golgi reassembly.

## Introduction

The AAA^+^ ATPases valosin-containing protein (VCP)/p97, known as cdc48 in *Saccharomyces cerevisiae*, is an essential and highly conserved protein involved in numerous cellular functions, such as regulating protein homeostasis^1^, endolysosomal trafficking^2^, membrane remodeling^3,4^, chromatin regulation^5^, autophagy^6–8^, Golgi formation^9^, and others^10,11^. Its core function is to unfold ubiquitylated proteins in an ATP dependent fashion^12–15^. VCP achieves its versatility to facilitate such diverse cellular functions by engaging many different cofactors^16^, including adapter proteins that bind to VCP and mediate its interactions with substrates and/or additional binding partners^17,18^. How adapter binding governs VCP function has been extensively studied in the context of its role in the ubiquitin proteasome system (UPS)^19,20^. Structural and biochemical studies have played a significant role in providing a detailed understanding of how VCP unfolds ubiquitylated substrates by capturing various states of this AAA^+^ ATPase, with and without binding partners and substrates^13,19^. Although these studies provided a comprehensive framework for understanding ATPase and unfoldase activity of VCP, less is known about how VCP in conjunction with multiple binding partners engages in activities outside the UPS. Specifically, how VCP cooperates with the deubiquitinating enzyme (DUB) valosin contain protein p97/p47 interacting protein 1 (VCPIP1) and, the adapter, p47 in Golgi reassembly, which does not seem to require substrate extraction from the membrane and/or its unfolding, remains poorly understood at the molecular level^21^.

Previous studies have established that post-mitotic Golgi reassembly requires presence of VCP, as well as two adapter proteins, VCPIP1^22^, and p47^23,24^. VCPIP1, also known as VCIP135, is part of the ovarian tumor protease (OTU) family of DUBs, an essential family of enzymes that remove ubiquitin tags from proteins to regulate protein homeostasis and function^25,26^. In Golgi, mono-ubiquitination of syntaxin-5 (Syn5), a t-SNARE protein essential for Golgi reassembly^27^, allows for the regulation of both assembly and disassembly processes during cell division. VCPIP1, in complex with VCP and p47, deubiquitinates Syn5 thereby controlling the affinity of the SNARE proteins^28^, allowing deubiquitinated Syn5 to bind to v-SNARE Bet1, resulting in membrane fusion and Golgi reassembly^22,23,29,30^.

To understand how VCP engages with VCPIP1 and p47 to mediate Golgi reassembly, we used cryogenic electron microscopy (cryo-EM) to determine high resolution structures of VCP-VCPIP1 and VCP-VCPIP1-p47 complexes capturing interaction states with high conformational flexibility. Our structures reveal that VCPIP1 exhibits a bivalent binding mode through two distinct regions of VCP. The VCPIP1 ubiquitin regulatory X, UBX, domain engages the N-domain of VCP, which is a well-established primary site of adapter protein binding^23^. Interestingly, we also identified a region on VCPIP1 that engages VCP at its C-terminal domain (D2)^31^. After biochemical validation of its functional relevance, we named this region “<u>VCP i</u>nteracting <u>d</u>omain” (VCPID). We found that while the UBX domain was essential for binding, the VCPID was important for DUB activity. We also determine the structure of the VCP-VCPIP1-p47 ternary complex, which reveals that the p47 UBX domain outcompetes the VCPIP1 UBX domain on the VCP N-domain, which may suggest dynamic competition for N-domain binding, as seen with other VCP cofactors^16,32,33^. Lastly, we show in cell-based studies that VCPIP1 UBX domain, VCPID and its DUB activity are all required for Golgi reassembly.

## Results

### VCP interacts with VCPIP1 through multivalent interactions

To reconstitute the VCP-VCPIP1 complex for structural studies, we co-expressed and purified full-length VCPIP1 and VCP (**Extended Data Fig. 1a**, see **Fig. 1a** for schematics of domain organization indicating domain boundaries) from Expi293 cells with no addition of ATP or ATP analogs (see **Online Methods**). Two-dimensional (2D) class averages from initial cryo-EM data sets revealed the typical hexameric structure of VCP with additional blurred density visible at the carboxy (C)-terminal domain of VCP, indicating significant conformational heterogeneity of this region (**Extended Data Fig. 1b**). To stabilize the complex, we used BS3 chemical crosslinking prior to grid preparation and cryo-EM data collection (**Extended Data Fig. 1c**). From this sample, initial 3D reconstructions revealed two binding sites of VCPIP1 on VCP (**Fig. 1b**). We obtained a consensus reconstruction of the VCP-VCPIP1 complex at an overall resolution of 2.3 Å (see **Online Methods** and **Extended Data Fig. 2** for details). To better resolve these regions, we used symmetry expansion followed by focused refinement, with masks focusing on the N- and C-terminal ends of VCP. This enabled us to obtain improved maps of specific subsections, including a 2.9 Å map resolving three VCPIDs bound to the D2 domains of VCP without symmetry expansion, a 2.9 Å map of one VCPID with stalk region bound to the bottom interface of VCP D2 dimer and a 3.1 Å map of the VCPIP1 UBX domain bound to the N-terminal domain of VCP, both with symmetry expansion (**Extended Data Fig. 2b −2e,** see **Table 1** for details).

**Figure 1:**
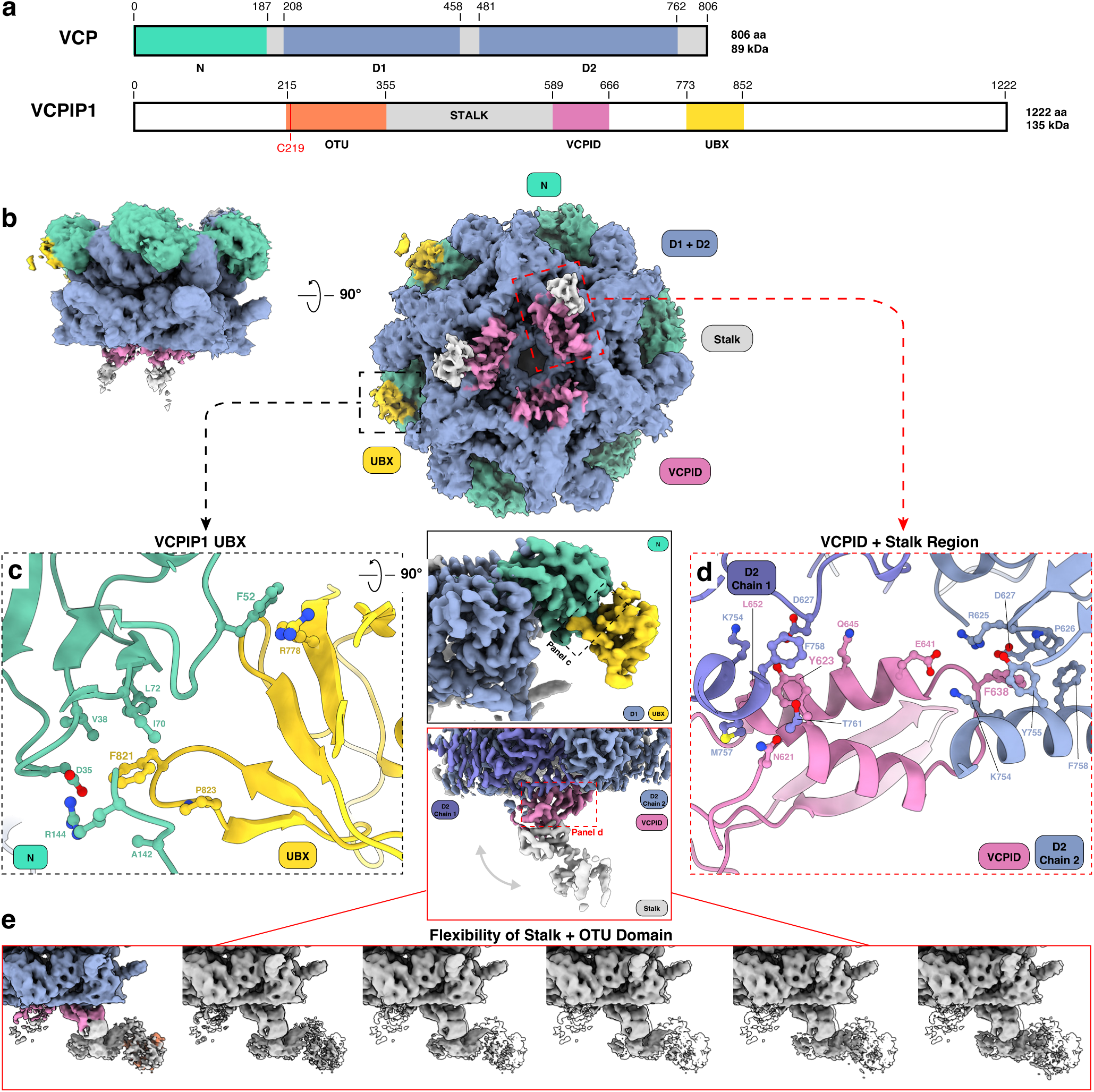
Cryo-EM structure depicting VCPIP1 as a multivalent binder of VCP. (a) Domain organization of VCP (top) indicating the three main domains: N-domain (green) and two ATPase domains (D1 and D2, blue), and VCPIP1 (bottom), indicating the OTU deubiquitinase (DUB) domain (orange) with the location of the catalytic cysteine (C219) (red), stalk region (grey), newly annotated VCPID (pink) and the UBX domain (yellow) (b) overall structure of the VCP- VCPIP1 complex in two orientations, color coded using the same colors as in (a), with two zoomed in regions focusing on two areas of VCP where additional density assigned to different regions of VCPIP1 are present (c) zoom in on the VCP N-domain interface with the VCPIP1 UBX domain displaying key interactions and features (d) zoom in on the VCP D2-domain dimer and its interaction with the newly defined VCPID (e) heterogeneity in our cryo-EM maps caused by the dynamic nature of VCPIP1 stalk and OTU domain (state 1 colored and then depicted in an outline, subsequent states in grey to highlight movement).

Of the two identified binding regions, we were able to clearly assign the VCPIP1 UBX domain (aa 773 – 852) and identified the larger interacting domain to the region spanning the stalk region (aa 356-588) and the domain of unknown function as VCPID (aa 589 – 666), which we newly characterize in this study **(Extended Data Fig. 1d)**. The VCPIP1 UBX domain is bound to the flexible N-domain of VCP limiting the resolution of the initial reconstructions which was addressed with C6 symmetry expansion to obtain a high-resolution map and build an atomic model (**Fig. 1c**). We observed that the VCPIP1 UBX domain folds into an archetypal UBX domain structure. The interaction interface is mainly formed by F821 on the UBX domain, buried in a hydrophobic pocket formed by VCP V38, I70, and L72 **(Extended Data Fig. 1e)**. Additional contacts are made between VCPIP1 P823 and VCP A142. We also identified a possible cation-π interaction between VCPIP1 UBX domain R778 and VCP F52 (**Fig. 1c**).

In addition to the anticipated VCPIP1 UBX domain interaction with the VCP N-domain, we observed density corresponding to three VCPIP1 interfacing with the D2 domain dimers of VCP (**Fig. 1b**), which we later assigned to the VCPID domain of VCPIP1. VCPID consists of a β-hairpin loop (aa 590-609) and two alpha helices (α1 (aa 622 - 633) and α2 (aa 640 – 654)) (**Fig. 1d, Extended Data Fig. 1d**) and was determined to have no domain conservation amongst the PDB^34,35^. The observed stoichiometry between VCP and VCPIP1 is 2:1 and each VCPIP1 VCPID contacts two D2 domains (D2 chain 1 and D2 chain 2, **Fig. 1b**). The VCPIP1 VCPID interacts with the dimer through its alpha helices, while the β-hairpin loop connects to the VCPIP1 stalk region (**Fig. 1d**). Notably, VCPID Y623 makes π-π interactions with F758 from VCP D2 chain 1 and engages through insertion in a hydrophobic pocket formed by D627, K754 and T761 on the same chain **(Extended Data Fig. 1f)**. VCPID N621, D627 and K754 also interact with VCP M757, L652 and Q645, respectively (**Fig. 1d**). The primary contacts of VCPID with the VCP D2 chain 2 are formed through insertion of VCPID F638 into a hydrophobic pocket formed by P626, Y755, and F758 (**Fig. 1d, Extended Data Fig. 1f**). This interface is further stabilized by a salt bridge formed between VCP K754 (D2 Chain 2) and VCPID E641 (**Fig. 1d**). The three VCPIDs interact exclusively with VCP and make no inter-domain contacts (**Extended Data Fig. 2**).

We observed variable quality of density for the VCPIDs, more specifically the β-hairpin loops, suggesting that there is compositional and/or conformational heterogeneity in the binding region. To enrich the dominant conformation, we again used focused classifications and refinements, and we were able to visualize additional density corresponding to the stalk region that binds to the VCPIP1 OTU domain (2.9 Å resolution; **Fig. 1b**). Further, this allowed us to visualize the flexible nature of the OTU domain (aa 215-355), which contains the catalytic cysteine (C219) responsible for DUB activity, and the stalk region that connects to VCPID (**Supplemental Movie 1**). The flexible nature of the stalk and the OTU domain prevented us from obtaining high resolution structure for these regions of VCPIP1, thus a portion of the stalk and all of the OTU domain are excluded from model building **(Extended Data Fig. 2**). Nonetheless, we analyzed different 3D classes to evaluate the movement of the stalk and the OTU domain. From the density visible, the stalk region moves ∼4 Å and the OTU domain moves ∼13.5 Å from the first to the last state (**Fig. 1e**).

Taken together, our cryo-EM structure of VCPIP1 bound to VCP revealed that VCPIP1 binds VCP in 1:2 stoichiometry. Each VCPIP1 molecule makes two distinct contacts with VCP, one mediated by the UBX domain that engages the N-domain of a VCP protomer, and one with the VCPID that forms contacts with D2 domains of two adjacent VCP protomers, enforcing the 1:2 stoichiometry. These interactions position the catalytic domain of VCPIP1 in close proximity to the end of the central pore of VCP. We also observed that the complex is highly dynamic, including within the N-domain of VCP and throughout VCPIP1.

### The UBX domain anchors VCPIP1 to VCP

Our structural analysis showed that both the UBX domain and VCPID domains of VCPIP1 bind VCP. To dissect their roles further, we measured binding affinities of wild type VCPIP1 (VCPIP1^WT^) and different VCPIP1 domain deletion constructs to VCP: VCPIP1^ΔVCPID^ (aa Δ589-666); VCPIP1^ΔUBX^ (aa Δ773-852); and VCPIP1^ΔVCPID ΔUBX^ (aa Δ589-666, Δ773-852) (**Fig. 2**). Using a time-resolved fluorescence resonance energy transfer (TR-FRET) assay, we observed that the two constructs with an intact UBX domain, VCPIP1^WT^ and VCPIP1^ΔVCPID^, bind to VCP with comparable affinity (K_D_ = 79.5 ± 7.0 nM and K_D_=67.9 ± 4.8 nM, respectively) (**Fig. 2a, 2b**). In contrast, the constructs lacking the UBX domain, VCPIP1^ΔUBX^ and VCPIP1^ΔVCPID ΔUBX^, did not show measurable affinity for VCP (**Fig. 2c, 2d**). These results reveal that the UBX domain of VCPIP1 is critical for VCP binding. This is in agreement with previous studies that identified the C-terminal region of VCPIP1, where the UBX domain is located, as critical for VCP binding^23^. Based on these studies, we propose that VCPIP1 UBX domain binding serves to anchor VCPIP1 to VCP, allowing VCPID to reach and engage the D2 domain region of VCP.

**Figure 2:**
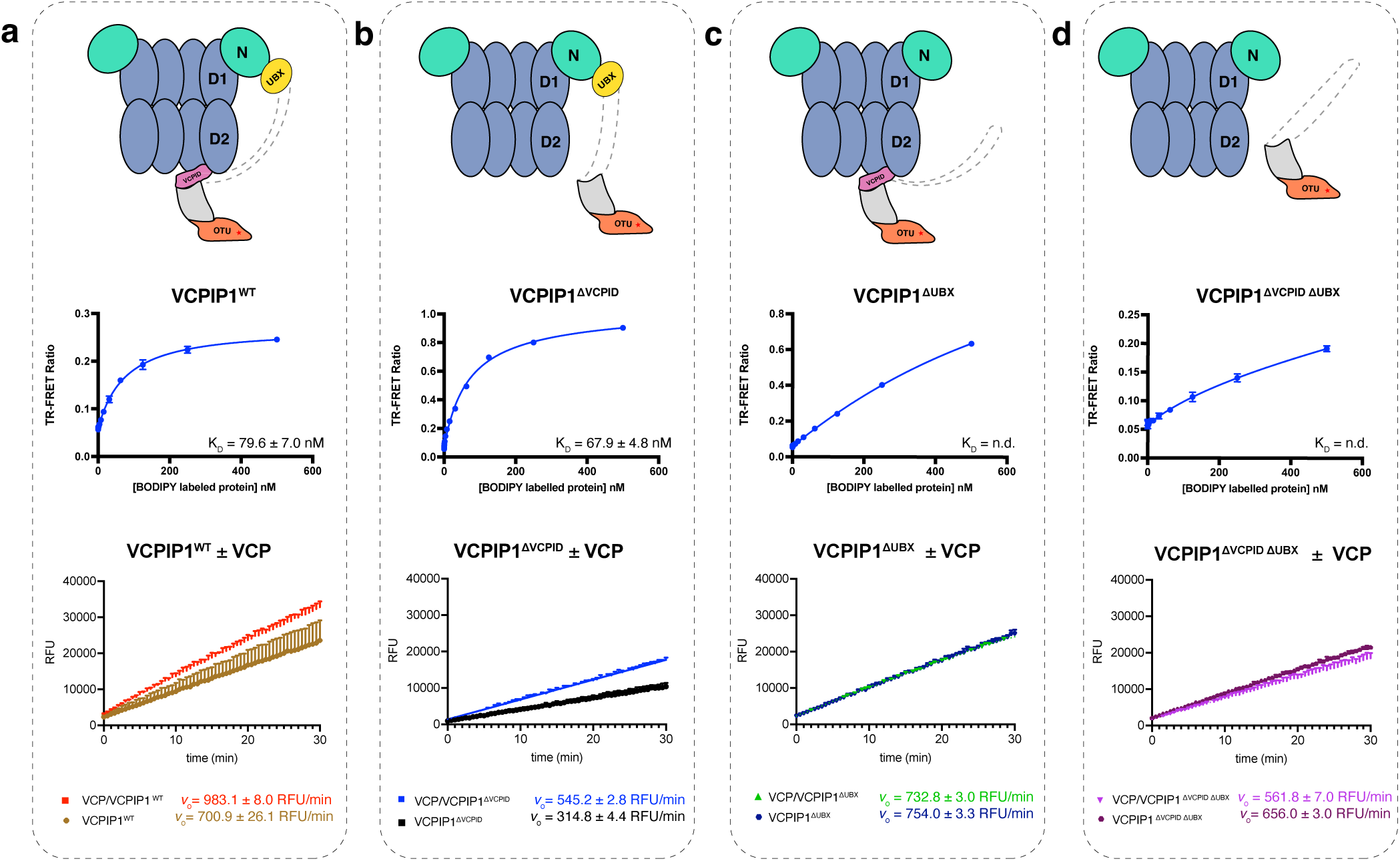
Characterization of VCPIP1 domain functionality in complex with VCP. First row shows cartoon depictions of VCP in complex with VCPIP1 containing various domain truncations, second row depicts TR-FRET binding results and last row depicts DUB activity results at 62.5 nM enzyme concentration. Dashed grey boxes are used to organize data with corresponding VCPIP1 truncation, (a) VCPIP1^WT^, (b) VCPIP1^ΔVCPID^, (c) VCPIP1^ΔUBX^ and (d) VCPIP1^ΔVCPID ΔUBX^, with the corresponding biochemical data.

### VCPID supports DUB activity of VCPIP1

We next asked whether complex formation with VCP influences the rate of deubiquitylation of substrates by VCPIP1. Using ubiquitin-rhodamine 110 (Ub-Rho) as a generic DUB substrate, we measured the initial velocities of the VCPIP1-mediated deubiquitylation reaction with and without full length VCP. In agreement with the literature^25,36^, we see that VCPIP1^WT^ is an active DUB (*v_0_* = 700.9 ± 26.1 RFU/min; **Fig. 2a, Extended Data Fig. 3a**). Removal of the UBX domain has no major effect on the initial velocity of the deubiquitylation reaction (*v_0_* (VCPIP1^ΔUBX^) = 754.0 ± 3.3 RFU/min; **Fig. 2c**). However, removing VCPID decreased the rate of ubiquitin hydrolysis significantly (*v_0_* (VCPIP1^ΔVCPID^) = 314.8 ± 4.4 RFU/min) (**Fig. 2b**). Surprisingly, the loss in activity was partially restored in the double deletion of UBX domain and VCPID (*v_0_* (VCPIP1 ^ΔVCPID ΔUBX^) = 656.0 ± 3.0 RFU/min; **Fig. 2d**), suggesting a potential auto-inhibitory mechanism involving the UBX. The presence of VCP increased the rate of deubiquitylation displayed by VCPIP1^WT^ (*v_0_* = 983.1 ± 8.0 RFU/min; **Fig. 2a**) and VCPIP1^ΔVCPID^ (*v_0_* = 545.2 ±2.8 RFU/min; **Fig. 2b**) but had no effect on the activity of VCPIP1^ΔUBX^ (*v_0_* = 732.8 ± 3.0 RFU/min; **Fig. 2c**) and VCPIP1^ΔVCPID ΔUBX^ (*v_0_* = 561.8 ± 7.0 RFU/min; **Fig. 2d**).

Collectively, these data suggest that binding to VCP enhances the rate of the deubiquitylation reaction catalyzed by VCPIP1^WT^ **(Extended Data Fig. 3b).** We observed that VCPID removal has the largest effect on the rate of the DUB activity, suggesting that this domain either helps in orienting the active site towards the substrate or allosterically activates DUB activity, both in the presence and absence of VCP **(Extended Data Fig. 3c)**. Importantly, removal of the UBX domain resulted in constructs that are not sensitive to the presence of VCP, which is in line with reduced affinity for VCP as determined using a TR-FRET assay (**Fig. 2a-2d**). Thus, for the highest level of VCP-dependent enhancement of deubiquitylation activity, both the UBX domain and the VCPID of VCPIP1 need to be present.

### Structure of the VCP-VCPIP1-p47 ternary complex

VCP activity in Golgi reassembly requires involvement of both VCPIP1 and p47^23^ (**Fig. 3a**). To provide a structural basis of how these two cofactors interact with VCP and a framework for understanding this process, we determined a cryo-EM structure of the VCP-VCPIP1-p47 complex **(Extended Data Fig 4a – 4c)**. After initial rounds of 3D classification, we were again able to observe density in the two binding regions on VCP, similar to what we observed for the VCP-VCPIP1 complex (see **Online Methods** and **Extended Data Fig. 5** for details). We used a combination of symmetry expansion and focused refinement to zoom in on these regions. We obtained a 3.1Å reconstruction revealing three VCPIDs bound at the C-terminal end of VCP without symmetry expansion, and a 4.0Å reconstruction of the N-domain region with symmetry expansion. The binding mode of the three VCPIDs bound to the D2 dimer is indistinguishable from the binding mode observed in the VCP-VCPIP1 complex, with the stalk regions connected to each VCPID clearly visible (**Fig. 3b**). Since the interactions between VCP and the VCPID in all structures are unchanged, we conclude that p47 binding does not affect how VCPIP1 engages the D2 domains of the VCP hexamer.

**Figure 3:**
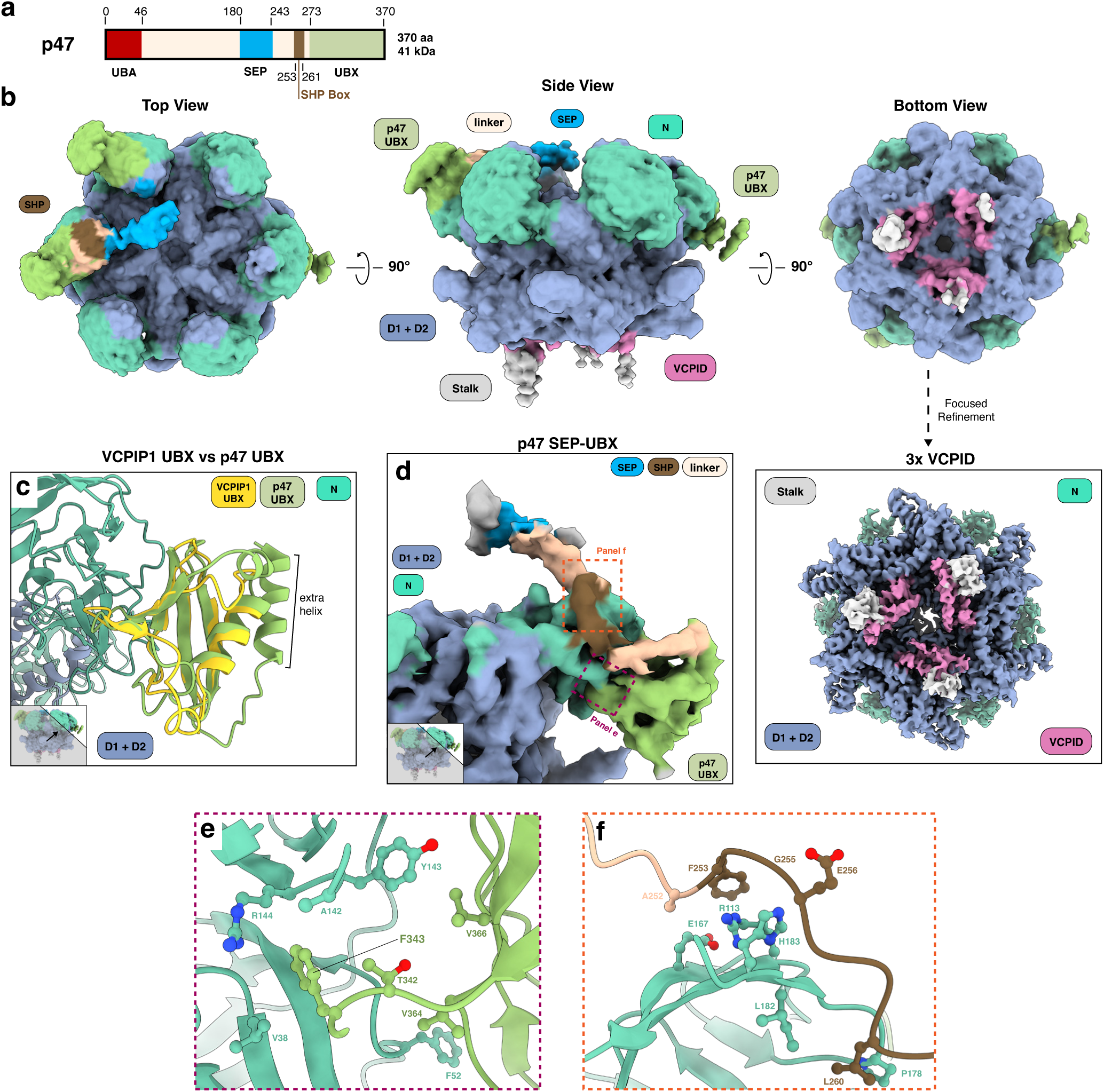
Cryo-EM structure of VCP bound to VCPID and adapter protein p47. (a) Domain organization of p47 showing the presence of its major domains: ubiquitin associated (UBA) (red), SEP domain (blue), SHP box (brown), UBX domain (lime green) and the connecting linker (beige); (b) low pass filtered (6.0 Å) map of VCP-VCPIP1-p47 ternary complex depicted in top, side and bottom views with a 90° rotation; (c) comparison of VCPIP1 UBX and p47 UBX domains; (d) zoom in on density for p47 bound to VCP N-domain; (e) zoom in of interactions formed by p47 UBX domain and VCP N-domain; (f) zoom in on interactions formed between p47 SHP binding site and linker region and VCP N-domain.

We next focused on the UBX domain observed on VCP N-domains. We find that there is clear density for three UBX domains, with two bound to N-domains in the up-position and one bound with N-domain in the down-conformation **(Extended Data Fig. 4d**, see **Online Methods, Fig. 3b**). Although the UBX domains from VCPIP1 and p47 share structural homology (**Fig. 3c**), the p47 UBX domain has an additional helix (aa 273-286), which allowed us to assign all three bound UBX domains as originating from p47 **(Extended Data Fig. 4e, 4f)**. Using AlphaFold3^37^ predictions of VCP N and D1 domain bound to p47, we were able to fit in the highest ranked predicted model into our density **(Extended Data Fig. 4g**, see **Online Methods)**. Compared to the VCPIP1 UBX domain, the p47 UBX domain binds to a partially overlapping area of the VCP N-domain (**Fig. 3d**). It also uses a phenylalanine, F343, to occupy a hydrophobic pocket within the N-domain defined by VCP’s V38, A142, Y143, and R144 (**Fig. 3e, Extended Data Fig. 4h**). Additional interactions are mediated by V364 and V366 of p47 UBX domain that engage with F42 and Y143 of the VCP N-domain, respectively (**Fig. 3e**). The extra helix of the p47 UBX domain is connected to a short loop (aa 244-272) which is connected to the SEP domain (aa 180-243) through a long linker that includes the SHP box (aa 253-261) (**Fig. 3d**), a motif previously reported as critical for VCP binding^38^. We observe significant, yet lower resolution, density for the linker to the SEP domain with the SEP domain potentially hovering over the central pore of VCP (**Fig. 3b**, see **Online Methods**). The SHP box acts a secondary binding site for p47 UBX domain as it has residues that interact with the top of the VCP N-domain, unlike VCPIP1 UBX domain (**Fig. 3d and 3f**). Specifically, L260 of p47 further stabilizes P178 and L182 on the VCP N-domain, G255 and E256 interact with H183 on VCP and F253 SHP approaches R113 and E167 (**Fig. 3f**). Lastly, there are additional contacts through A252 right next to the SHP motif, which interacts with E167 on VCP (**Fig. 3f**).

To measure and compare the affinities of VCPIP1 and p47 for VCP, we developed a TR-FRET displacement assay **(Extended Data Fig. 4i**). Unlabeled p47 and VCPIP1^WT^ were used to compete out a pre-formed complex of terbium labeled VCP and BODIPY-VCPIP1^WT^. We find that p47 (IC_50_ = 20.4 ± 2.8 nM) has a slightly higher affinity for VCP than VCPIP1^WT^ (IC_50_ = 74.54 ± 13.9 nM). Since p47 has a secondary binding site on the N-domain of VCP, compared to VCPIP1 UBX domain, this aligns well with the p47 UBX domain being exclusively observed in the VCP-VCPIP1-p47 complex structure. However, given that the apparent affinities are similar, that p47 and VCPIP1 UBX domains bind to overlapping locations on VCP, and that VCPIP1 UBX domain is the anchor point for VCP engagement based on our data (**Fig. 2**), it is likely that both p47 UBX domain and VCPIP1 UBX domain bind to VCP in a dynamic manner, which may be further modulated in the presence of substrate. Taken together, the cryo-EM structure of VCP-VCPIP1-p47 complex reveals how VCP is able to engage with p47 and VCPIP1 simultaneously and maps key points of interaction between VCP and the two adapter proteins involved in mediating Golgi reassembly. The binding appears to be dynamic and not mutually exclusive, suggesting multiple regulatory mechanisms.

### VCPIP1 catalytic activity and binding to VCP are required for Golgi reassembly

Next, we asked whether the interaction of VCPIP1 with VCP is necessary for the role of VCPIP1 in Golgi reassembly and assessed the involvement of different VCPIP1 domains. We developed a cellular system in which the knock-out of VCPIP1 resulted in a Golgi fragmentation phenotype that would be rescued by add-back of exogenous VCPIP1. We first used CRISPR/Cas9 to engineer A2058 cells with bi-allelic knock-out (KO) of VCPIP1. To observe Golgi bodies and thus Golgi reassembly, we stained cells with nuclear stain (DAPI) and a Golgi body stain (Golgi ID kit). Compared to wild type A2058 cells, VCPIP1 KO cells exhibited more fragmented Golgi as observed by confocal microscopy using maximal intensity projection image of the Z-stacks (**Fig. 4a** VCPIP1 WT vs VCPIP1 KO and **methods**). Next, the A2058 VCPIP1 KO cells were transiently transfected with either VCPIP1 wild type or mutant constructs (VCPIP1^ΔVCPID^, VCPIP1^ΔUBX^, VCPIP1^ΔVCPID ΔUBX^, and VCPIP1^C219A^) (**Extended Data Fig. 6a**). We observe more pronounced Golgi fragmentation in VCPIP1^KO^, VCPIP1^ΔVCPID^, VCPIP1^ΔUBX^, VCPIP1^ΔVCPID ΔUBX^ and VCPIP1^C219A^ conditions, as indicated by an increase in the number of maximal intensity spots and disruption of Golgi as seen by the gaps in the cell cytoplasm (**Fig. 4**). In contrast, transfecting the VCPIP1^WT^ construct into VCPIP1 KO cells (VCPIP1 rescue) phenocopied features of the WT VCPIP1 phenotype indicated by the low frequency of maximal intensity spots and the intact appearance of the cellular membrane (**Fig 4**). These studies suggest that VCPID, UBX domain and the catalytic cysteine are all essential for VCPIP1 activity in Golgi reassembly.

**Figure 4:**
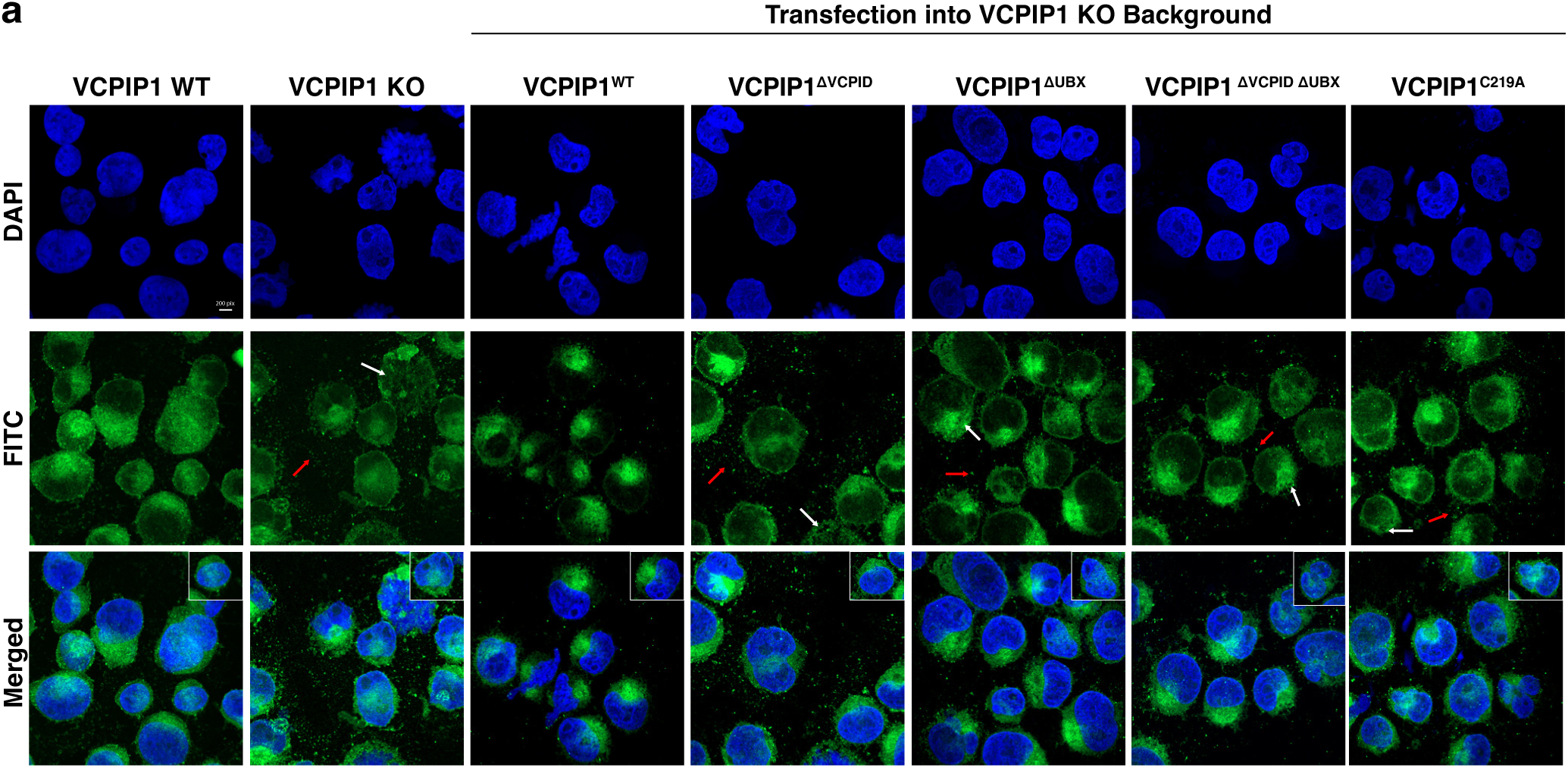
VCPIP1 and its interaction with VCP is necessary for Golgi reassembly. (a) Representative fluorescent images of A2058 WT and KO cells in which various VCPIP1 constructs were transiently transfected into the cells. Cells were stained with DAPI (first row) and GOLGI ID stain (second row). White arrows indicate “gaps” in the cytoplasm of the cell and red arrows depict maximal intensity spots. DAPI and FITC channels were merged as depicted in the third row. Representative cells from the merged image are highlighted on the top right of each merged image. Scale bar: 200 pixels.

## Discussion

VCP engages with a wide range of cofactor proteins to function in distinct cellular processes. However, our understanding of how VCP engages with different adapters in pathways outside the UPS remains poorly understood, especially in the context of its function in post-mitotic Golgi membrane biogenesis. Here, we provide structural insights into how VCP engages VCPIP1 and p47, two adapters that are critically involved in this process. Our structure of the VCP-VCPIP1 complex reveals that, unlike other VCP-adapter structures that show a single point of contact, VCPIP1 forms two distinct interfaces with VCP. This bivalent interaction is mediated by the well-characterized UBX domain bound to the N-terminus of VCP, and the newly characterized VCPID bound to the C-terminal VCP D2 domain. We observed a 2:1 VCP/VCPIP1 stoichiometry, with each UBX domain engaging a single N-domain of VCP, and each VCPID engaging two D2 domains from neighboring VCP protomers. Due to the highly flexible nature of this complex, we were unable to visualize the OTU domain and the position of the catalytic cysteine on VCPIP1; however, we captured VCPID and the stalk region that connects VCPID to the OTU domain, as well as observed the movement of the stalk. The bivalent mode of binding between VCP and VCPIP1 described here was also recently observed by *Vostal et al.*^39^. Furthermore, the interface between VCP D2 domains and VCPIP1 region we now annotate as VCPID was also recently reported by *Liao et al.*^40^. These two studies, which came out during the preparation of our manuscript, strongly support our conclusions about how VCPIP1 engages VCP.

Our binding and activity studies showed that, while VCPIP1 UBX domain serves as an anchor for VCP binding, VCPID is required for VCP-mediated enhancement of VCPIP1 DUB activity. Phenotypically, we qualitatively observe increased Golgi membrane fragmentation with VCPIP1^ΔVCPID^, VCPIP1^ΔUBX^, VCPIP1^ΔVCPIDΔUBX^ and VCPIP1^C219A^ suggesting that these regions and the DUB activity all have an important role in VCPIP1 function and its role in post-mitotic Golgi reassembly. Moreover, the VCP-VCPIP1-p47 structure provides context on how both VCPIP1 and p47 simultaneously bind to VCP, providing the first structural glimpse into the dynamic ternary complex that is crucial for Golgi reassembly.

Previous studies have demonstrated that VCP, in complex with VCPIP1, is necessary for cisternal regrowth^22^, and that p47 may serve to recruit the VCP-VCPIP1 to the substrate^24,28,32^. In that model, mono-ubiquitinated Syn5 is recognized by the UBA domain of p47^24,28^, which brings the substrate into the vicinity of VCPIP1 for deubiquitylation. Following deubiquitylation, Syn5 is free to bind Bet1 and form a SNARE complex resulting in membrane fusion, whereas VCPIP1 dissociates from VCP-p47, potentially driven by ATP hydrolysis induced conformational changes in VCP^28^. Therefore, based on this mechanistic proposal and our structural data, in the context of post-mitotic Golgi reassembly VCP-VCPIP1-p47 complex behaves as a multicomponent DUB, rather than the classical VCP ATPase/unfoldase machinery. Additional studies will be needed to test this mechanistic model further.

In conclusion, our studies indicate high degree of dynamics and flexibility in VCP-VCPIP1 and VCP-VCPIP1-p47 complexes and reveal a bivalent mode of engagement between VCP and VCPIP1, including the unique interface between the newly annotated VCPID of VCPIP1 and D2 domains from two VCP protomers. The exact consequence of this mode of binding remains to be determined; however, given that these interactions are maintained in the context of VCP-VCPIP1-p47 complex, we expect that they play a key role in ensuring efficient Syn5 deubiquitylation.

## Acknowledgements

This work was supported by NIH NCI 5F31CA281197-02, NIH NCI R01CA262188 (to E.S.F.) and NIH NCI R01CA233800 (to S.J.B.). We thank the staff at the Harvard Cryo-EM Center for Structural Biology for their outstanding support during grid screening and data collection and the staff at the Dana Farber Cancer Institute Molecular Imaging Core (MIC) for their microscopy training and support. We acknowledge SBGrid for assistance with software and high-performance computing^41^. We acknowledge Milka Kostic for her valuable input and critical feedback on the manuscript and members of the Buhrlage and Fischer labs for discussions, help and feedback.

## Author contributions

B.S., M.H., S.J.B., and E.S.F. conceived study and designed research plan. B.S. cloned and purified proteins, performed biochemical assays and cellular assays and, with the support of M.H., conducted cryo-EM structure determination. A.B. created the VCPIP1 KO cell line. H.Y. and I.J.M. provided technical insight and initial analysis for biochemical assays. B.S., M.H., S.J.B., and E.S.F. designed experiments, and B.S. analyzed and interpreted data. S.J.B. and E.S.F. supervised the study and acquired funding. B.S. prepared figures and wrote the original manuscript. M.H., S.J.B., E.S.F. reviewed and revised the manuscript. All authors approved of the final version of the manuscript.

## Extended Data Figures

**Extended Data Figure 1: Structural characterization of VCP-VCPIP1 complex.** (a) SDS-PAGE of purification of co-expressed VCP and VCPIP1 from Expi293 cells. (b) 2D classes from initial Talos Arctica session of the VCP-VCPIP1 complex without BS3 crosslinking. White arrows demonstrate VCPIP1 VCPID signal at the C-terminus of VCP. (c) SDS-PAGE of SEC fractions of BS3 crosslinked VCP-VCPIP1 complex that was concentrated for Titan Krios data collection. (d) Ribbon depiction of VCPIP1 VCPID. (e-f) Surface depiction of hydrophobicity found at the interaction sites for VCPIP1 UBX domain and VCPID, respectively. Hydrophobic areas are represented by surface mesh in yellow and pink, respectively.

**Extended Data Figure 2: Cryo-EM processing workflow for the VCP-VCPIP1 complex.** (a) Overview of processing workflow from raw micrographs to final maps. All steps were completed in cryoSPARC. All resolutions are given after post-processing. All colored volumes indicate selected classes of particles used for subsequent steps. (b-e) FSC plots (left top), 3DFSC plot (left middle), model-to-map FSC (left bottom), viewing direction distribution (right top) and main map colored according to local resolution (right bottom). (b) consensus EMD-XXXXX (c) 3xVCPID EMD-XXXXX (d) VCPID + stalk EMD-XXXX (e) VCPIP1 UBX domain EMD-XXXXX

**Extended Data Figure 3: DUB activity assay using ubiquitin rhodamine 110 (Ub-Rho) as substrate.** (a) Catalytic activity of VCPIP1^WT^ at different enzymatic concentrations and equal substrate (Ub-Rho) concentrations (500 nM). (b-c) Ub-Rho assay data reorganized with (b) and without (c) VCP at 62.5 nM enzyme concentration. (b) *v_0_* (VCP) = 3.809 ± 0.24 RFU/min. All other values are depicted in Figure 2.

**Extended Data Figure 4: Structural characterization of VCP-VCPIP1-p47 complex.** (a) SDS-PAGE gels depicting independently purified VCP (left), VCPIP1 (middle) and p47 (right). (b) SEC run on Superose6 Increase 10/300 GL (Cytiva) demonstrating traces of both VCP-p47 and VCP-VCPIP1-p47 complex formation. SDS-PAGE gel represents SEC fractions from VCP-VCPIP1-p47 complex formation, and the fractions concentrated for cryo-EM analysis on the Titan Krios. (c) 2D class averages of the VCP-VCPIP1-p47 complex from Titan Krios data set. (d) Classes from heterogeneous refinement job (highlighted in dashed green boxes in **Extended Data Fig. 5**) with N-domains of VCP in the down-conformation (top) and up-conformation (bottom). (d) VCPIP1 (yellow) and p47 UBX (lime green) domains individually fit in p47 UBX domain density (lime green, transparent). (f) VCPIP1 (yellow) and p47 UBX (lime green) domain alignment highlighting the extra helix on p47. (g) Highest rank prediction of p47 interaction with VCP N and D1 domains depicted by the resulting PAE plot (left) and model fit into density (right) (h) Surface depiction of hydrophobicity found at the interaction site for the p47 UBX domain and VCP N-domain represented similar to **Extended Data Fig. 1**. (i) Schematic and results of TR-FRET displacement assay of unlabeled VCPIP1 (red) and p47 (blue) displacing BODIPY-VCPIP1 from Tb-VCP.

**Extended Data Figure 5: Cryo-EM processing workflow for the VCP-VCPIP1-p47 complex.** (a) Overview of processing workflow from raw micrographs to final maps. All steps were completed in cryoSPARC. All resolutions are given after post-processing. All colored volumes indicate selected classes of particles used for subsequent steps. (b-c) FSC plots (left top), 3DFSC plot (left middle), model-to-map FSC (left bottom), viewing direction distribution (right top) and main map colored according to local resolution (right bottom). (b) 3xVCPID EMD-XXXXX (c) p47 EMD-XXXXX

**Extended Data Figure 6: Analysis of Golgi reassembly with VCPIP1 domain truncations.** (a) Western blot, blotting for VCPIP1, demonstrating successful transient transfection of VCPIP1 constructs in A2058 VCPIP1 KO cells.

**Extended Data Figure 7: Uncropped SDS-PAGE and western blots.** All original blots used in the extended data figures with cropped regions marked (black box).

**Supplementary Movie 1: Range of motion of VCPIP1 bound at the C-terminal of VCP.** 3D variability analysis demonstrating that the stalk and OTU domains of VCPIP1 are highly flexible at the bottom of VCP.

## Materials and Methods

### Cloning, protein expression and purification

Full-length VCPIP1 in pDEST_CMV with a N-terminal Strep tag was acquired from Wade Harper ^42^ and VCP was a gift from Nico Dantuma (VCP in pEGFP-N1 [Addgene plasmid #23971]). From there, the VCP coding sequence was moved into a modified pDARMO (pDarmo.CMVT_v1 was a gift from David Sabatini [Addgene plasmid #133072]) expression plasmid with a N-terminal FLAG tag. Mutant and truncated VCPIP1 constructs were generated by excising and replacing sequences using primers with PCR, T4 polynucleotide kinase ligation (mixture of DNA, T4 ligation buffer, PNK (NEB M0201S), T4 DNA ligase (NEB, M0202S) and H_2_0) and 1 hour incubation at 37°C^43^.

All VCPIP1 variants and VCP were expressed transiently in Expi293 (Thermo Fischer Scientific, A14635) following the manufacturer’s manual. Cells were harvested 60-65 hours post-transfection and lysed by sonication in lysis buffer (50 mM HEPES pH 7.4, 200 mM NaCl, 5% glycerol) supplemented with protease inhibitors. The lysate was cleared by ultracentrifugation (39,000 rpm, 45 minutes) and then incubated with either StrepTactin®XT 4Flow high-capacity resin (IBA life sciences, 2-5030-002) or FLAG-antibody-coated beads (Genscript, L00432). Bound proteins were eluted with 50 mM of biotin or 0.2 mg/mL 1xFLAG (DYKDDDDK) peptide. Proteins were further purified by ion exchange chromatography (IEX) with a Poros 50HQ (Thermo Fisher Scientific, 1255911) column, eluting with a linear NaCl gradient from 200 - 750 mM. Elution fractions were concentrated and polished by size exclusion chromatography (SEC) in SEC buffer (30 mM HEPES pH 7.4, 150 mM NaCl) using Superose6 Increase 10/300 GL (VCP alone) and Superdex200Inc 10/300 GL (VCPIP1 alone) columns (Cytiva). For purification of co-expressed VCP-VCPIP1 complex, ion exchange chromatography was omitted.

p47 was cloned into pNIC-Bio2 with an N-terminal His-6-TEV tag. For expression, LOBSTR *E. coli* expression strains (Kerafast, EC1002) were transformed, and a 1 L culture was grown in TB at 37 °C to OD_600_=0.6, and induced with 0.5 mM Isopropyl β-D-1-thiogalactopyranoside (IPTG). After induction, temperature was decreased to 18 °C and the protein was expressed overnight. The culture was centrifuged (4,000 rpm, 20 minutes), resuspended in lysis buffer (50 mM HEPES pH 7.4, 200 mM NaCl, 20 mM imidazole, 5% glycerol, benzonase and protease inhibitors), lysed using sonication, and the lysate was cleared using ultracentrifugation (39,000 rpm, 60 minutes). Clarified lysate was applied to high affinity Ni-charged resin (Genscript, L00223) and eluted with increasing imidazole concentrations (20-750 mM). p47 was further purified using ion exchange chromatography – the fractions were diluted to 50 mM NaCl, applied to a PorosHQ and eluted with a NaCl gradient from 50-750 mM NaCl. Peak fractions were concentrated using centrifugal concentrators (30 kDa MWCO, Millipore Amicon, UFC3090) and applied to a Superdex75 Increase 10/300 GL (Cytiva) equilibrated in SEC buffer (30 mM HEPES pH 7.4, 150 mM NaCl). Final protein samples of all constructs were flash-frozen in liquid nitrogen and stored in −80 °C.

### EM sample preparation and data collection

For data set 1 (VCP-VCPIP1): VCP-VCPIP1 was crosslinked with a 400x molar excess of bis(sulfosuccinimidyl)suberate (BS3) crosslinker (9.8 µM BS3 (12 mM), 0.5 mg VCP-VCPIP1 sample (5.2 mg/mL) and SEC buffer), quenched with 100 mM Tris and loaded on Superose6 Increase 10/300 GL (Cytiva) which was equilibrated with SEC buffer for further purification. Final concentrated sample (2.4 mg/mL) was diluted to 1.8 mg/mL and mixed with CHAPSO (0.2 mM final concentration) directly prior to grid preparation. 4 µL of sample were applied to the grid and blotted for 3 s followed by a 3 s post-blot incubation before vitrification.

For data set 2 (VCP-VCPIP1-p47): Individually purified VCPIP1, VCP and p47 were mixed and incubated for 30 minutes on ice, loaded on Superose6 Increase 10/300 GL (Cytiva) and concentrated (30 kDa MWCO) to 4.7 mg/mL. Sample was diluted to 2.0 mg/mL and CHAPSO (0.2 mM final) was added directly prior to grid preparation. 4 µL of sample was applied once to the grid and blotted for 3 s followed by a 3 s post-blot incubation before vitrification.

Quantifoil 1.2/1.3 300, N1-C14Cu30-50 grids were glow discharged for both data sets at 20 mA, 60 s, 39 Pa. Subsequently, grids were vitrified in a Leica EM-GP operated at 10 °C and 90% relative humidity with 10 s pre-blot time.

Data Set 1 was collected using SerialEM^44^ (v3.8.6) in a Thermo Scientific Titan Krios equipped with a Gatan Quantum image filter (20 eV slit width) and a post-GIF Gatan K3 direct electron detector. Movies were acquired at 300 kV at a nominal magnification of 105,000x in counting mode. 11,590 movies (50 frames each) were recorded with 3 exposures per hole and 9 holes per stage position resulting in 27 image acquisition groups. Defocus was varied from −0.9 - −2.2 µm and total dose, pixel size and exposure time were 53.4 e–/ Å^2^, 0.83 Å and 2.83 s, respectively.

Data Set 2 was collected using EPU (v3.7) in a Thermo Scientific Titan Krios equipped with a Selectrix energy filter (10 eV slit width) and a post-GIF Falcon4 direct electron detector. Movies were acquired at 300 kV at a nominal magnification of 165,000x in counting mode. 12,261 movies (49 frames each) were recorded with (3) per hole. Holes per stage position were calculated by EPU software. Defocus was varied from −0.8 - −2.2 µm and total dose, pixel size and exposure time were 49.2 e–/ Å^2^, 0.73Å and 2.87 s, respectively.

### Data processing and model building

Initial processing was done with cryoSPARC^45^ (v3.3.1) and later processing and refinement was done with cryoSPARC (v4.6.0). Resolutions are stated based on the Fourier shell correlation (FSC) 0.143 threshold criterion^46^.

Data set 1 (VCP-VCPIP1): 11,580 movies were corrected for beam induced motion and CTF was estimated on-the-fly using cryoSPARC live. A total of 2,050,270 particles were extracted (1.13 Å/pix) after crYOLO^47^ particle picking from 9,365 curated micrographs. The consensus structure was resolved at 2.3 Å using homogenous refinement after two rounds of heterogeneous refinement. 3D variability analysis was performed on these particles (1,369,446), followed by homogenous refinement, leading to a 2.9 Å reconstruction (107,859 particles) of three VCPID regions binding to the C-terminal end of VCP. To resolve more density for one VCPID, C6 symmetry expansion was done on the particles from the consensus refinement resulting in 8,216,676 particles followed by local clustering using 3D variability. A 2.9 Å reconstruction was resolved after local refinement of 592,039 particles. Lastly, the symmetry expanded particles were downsampled (2.6 Å/pix) followed by local clustering with 3D variability, 3D classification, 3D variability, and local refinement to resolve density for VCPIP1 UBX domain at a resolution of 3.1 Å. Final particle stacks were polished using reference based motion correction (restoring the particles to 0.83 Å/pix) and final refinements were performed with per-particle defocus estimation and correction of higher order CTF aberrations per-acquisition group **(Extended Data Fig. 2**). All final maps were postprocessed with deepEMhancer^48^, and these maps, together with the sharpened and unsharpened maps from cryoSPARC were used for model building. Initial models for VCPIP1 consensus, 3xVCPID, VCPID, and VCPIP1 UBX domain were generated by rigid-body docking (in ChimeraX (v 1.6)^49^) in a combination of VCP from the Protein Data Bank (PDB) (PDB:5FTK^14^) with truncations of the AlphaFold model for VCPIP1 (UBX domain, VCPID and stalk region) (AF-Q96JH7-F1-v4). The combined models were first flexibly fit using ISOLDE^50^, followed by residue-by-residue modeling and inspection with COOT (v 0.9.8)^51^ and then finally refined using phenix.real_space_refine^52^ using reference restraints, after preparation using phenix.ready_set^53^.

Data set 2 (VCP-VCPIP1-p47): 12,261 movies were corrected for beam induced motion and CTF was estimated on-the-fly using cryoSPARC live. A total of 2,003,371 particles were extracted (at 1.96 Å/pix) from 12,129 curated micrographs after cryoSPARC template picking. Two rounds of heterogenous refinement were done initially to classify out a VCP dodecamer class. Two subsequent rounds of heterogenous refinement were done to classify out VCP states. From the fifth heterogenous refinement run, one class that had obvious VCPID density was used to resolve the 3xVCPID structure of three VCPID binding to the C-terminal end of VCP. For the 3xVCPID structure, homogeneous refinement was done after heterogenous refinement and further classification was carried out with 3D variability. Two classes from the 10 classes of 3D variability were selected for local refinement, followed by two rounds of heterogenous refinement. Lastly, local refinement was done resulting in a 3.1 Å reconstruction from 77,321 particles. For the p47 UBX domain structure, particles from heterogeneous refinement were combined and C6 symmetry expansion was applied, resulting in 4,293,558 particles. Local clustering with 3D variability was carried out with a mask on the UBX region, followed by 3D classification, another round of 3D variability, local refinement, back-to-back rounds of 3D variability and a final local refinement, resulting in a 4.0 Å reconstruction **(Extended Data Fig. 5**). The model from data set 1 was used as a starting point for model building, and AlphaFold3^37^ was used to predict the p47 linker, the SHP motif and the SEP domain interaction with VCP. The highest rank prediction from AlphaFold3 of VCP N and D1 domains and full length p47 was used as the initial model for p47 density bound to VCP N-domain. This model was first flexibly fit using target restraints in ISOLDE, refined using phenix.real_space_refine with reference restraints, and finally inspected with COOT. Local resolution ranges are given based on 0-75% percentile in local resolution histograms^54^. Structural biology applications used in this study were configured by SBGrid^41^.

### Interface Residues

Initial identification of interacting residues between VCP and VCPIP1 was determined based on visual inspection of density in parallel to scores corresponding to interaction using the script: residue_energy_breakdown_script.sh. We also used PDBePISA^55^ to confirm residues with the highest buried area percentage. Residue predictions for p47 and VCP N-domain interaction are based on the AlphaFold3 model and were similarly cross-validated using PDBePISA and the script.

### Fluorescent protein labeling for TR-FRET assays

VCPIP1 constructs (VCPIP1^WT^, VCPIP1^ΔVCPID^, VCPIP1^ΔUBX^, VCPIP1^ΔVCPID ΔUBX^) and p47 were labeled with BODIPY and VCP was labeled with Terbium (Tb) (CoraFluor : R&D systems, Cat. No. 7920) in a 1:1 molar ratio as previously described^56^. The samples were incubated for 1 hour at room temperature. After incubation, the samples were quenched with 20 mM Tris pH 8, incubated for 10 minutes and spun in a Zeba Spin Desalting Columns (Thermo Fisher Scientific, 89882) to remove excess BODIPY and Tb and buffer exchange back into SEC buffer. A280 and A503 readings were taken to calculate degree of labeling (DOL) for BODIPY labeled proteins; VCPIP1^WT^ (0.23), VCPIP1^ΔVCPID^ (DOL=0.62), VCPIP1^ΔUBX^ (DOL = 0.54), VCPIP1^ΔVCPID ΔUBX^ (DOL = 0.44), p47 (DOL =0.27). The extinction coefficient of 80,000 M^-1^cm^-1^ was used for BODIPY at 503 nm. A280 and A340 readings were taken to calculate the DOL of Tb-VCP (DOL =0.33). The extinction coefficient at A_340_ of CoraFluor was equal to 22,000 M^-1^cm^-1^ as described^56^.

### TR-FRET binding and competition assays

For binding assays, Tb-VCP was mixed with TR-FRET buffer (25 mM HEPES pH 7.4, 150 mM NaCl, 0.1% BSA, 0.05% Tween-20). A volume of 7.5 µL of the mixture was dispensed into each well of a 384-well plate (Corning, 4514). The serial dilution of labelled protein was performed in 96-well plate, and 7.5 µl of the serial dilution mixture was added to each well of the 384-well plate. The final assay volume per well was 15 µL. The plate was incubated for 1 hour at room temperature. Fluorescence signals were measured using a PheraStar FS plate reader (BMG Labtech). Tb was excited at 337 nm and emitted at 490 nm (Tb) and 520 (BODIPY) which were recorded with a 70 µs delay over a window of 600 µs. The plates were measured for 6 cycles. All assays were performed as technical triplicates. Data analysis was done using nonlinear regression in GraphPad Prism (v10.2.3).

For displacement assays, demonstrating the displacement of BODIPY-VCPIP1^WT^ by unlabeled-VCPIP1^WT^ or p47 from Tb-VCP, 10 nM BODIPY-VCPIP1^WT^ was incubated with 10 nM Tb-VCP. Serial dilution of unlabeled proteins (VCPIP1^WT^ or p47) was prepared in a 96-well plate. All other conditions to the assay were performed the same as the binding assays described above.

### Ubiquitin Rhodamine Assay

For Ub-Rho activity assays, enzyme concentration varied while substrate (Ub-Rho) concentration remained constant. Each VCPIP1 construct (VCPIP1^WT^, VCPIP1^ΔVCPID^, VCPIP1^ΔUBX^, VCPIP1^ΔVCPID ΔUBX^) was tested with and without VCP. VCP was also tested alone as a control. Enzyme dilutions were prepared in assay buffer (50 mM Tris pH 8.0, 0.5 mM EDTA, 5 mM TCEP, 11 µM ovalbumin), for final concentrations ranging from 500 nM to 7.8 nM. 10 µL of enzyme dilutions were distributed in a 384-well plate (Corning, 3820). Ub-Rho solution was prepared in the same assay buffer, for a final concentration of 500 nM. 10 µL of Ub-Rho solution was transferred to the 384-well plate and quickly centrifuged before reading on the CLARIOstar (BMG Labtech). Fluorescence was measured for 1 hour, every 30 seconds at an excitation and emission of 487 nm and 535 nm, respectively. Fluorescence over time was plotted and initial velocities analyzed using simple linear regression in GraphPad Prism. All assays were performed in duplicate.

### CRISPR CAS-9 KO of VCPIP1

#### Cell Culture

A2058 (a gift of Peter Sorger’s lab) and HEK-293T (ATCC catalog # CRL-3216), were cultured and maintained in DMEM (Gibco, 11965092) supplemented with 10% (v/v) fetal bovine serum (Sigma, F0926) and 100U/mL penicillin + 0.1mg/ml streptomycin (Gibco, 15070063). Cells were grown and maintained in tissue-culture treated dishes (Corning, CLS430167) at 37 °C with 5% CO_2_ in a water-jacketed CO_2_ incubator. For passaging, cells were washed with PBS (Gibco, 10010023) detached using 0.25% trypsin/EDTA (Gibco 25200056) for passaging. All cell lines were routinely tested for mycoplasma and verified to be mycoplasma free (Lonza, LT07-703).

#### Cas9 Expressing Cell Lines

A2058 were transduced with Cas9-Flag via lentiviral infection (Cas9-Flag plasmid was a gift of Peter Sorger’s lab). Following blasticidin selection (Gibco, A1113903), a sample of cells were harvested, pelleted by centrifugation (5,000 x g for 10 minutes at 4 °C), lysed on ice with lysis buffer (20 mM Tris pH 8, 150 mM NaCl, 1% NP-40, 10% glycerol, 1 mM TCEP and protease inhibitor cocktail (Thermo Scientific, 78429) and clarified by centrifugation after 30 minutes. Protein content was quantified by BCA (Thermo Scientific, 23225). Lysate was diluted to 2 mg/mL in lysis buffer. 4x LDS sample buffer (Thermo Fisher, B0008) supplemented with 10% BME was added to each sample. Following mixing, samples were heated to 95 °C for 10 minutes. Then samples were resolved by SDS-PAGE and analyzed by Western blot to confirm FLAG expression (Invitrogen, MA1-91878) with GAPDH control (Cell Signaling Technology, 2118)

#### Creation of stable VCPIP1 KO Cell Line

A2058 VCPIP1 KO cells were generated using the Dharmacon Edit-R crRNA system. Lyophilized crRNA (Horizon Discovery; VCPIP1 #1 CM-019137-01-0002; #2 CM-019137-02-0002; CM-019137-03-0002; #4 CM-019137-04-0002; NTC #1) and tracrRNA (Horizon Discovery, U-002005-05) were resuspended in buffer (Horizon Discovery, B-006000-100) to a final concentration of 10 uM per manufacturers protocol. A2098-Cas9-Flag cells (40,000 cells) in media without antibiotic were seeded in 12-well plates (Corning, 3513) and allowed to adhere overnight. Mixtures of tracrRNA and all 4 VCPIP1 guides, tracrRNA and NTC guide, and Dharmafect 1 (Horizon Discovery, T-2001-01) were prepared in Opti-MEM (Gibco, 31985070). Dharmafect mixture was added in equal volumes to the tracrRNA-VCPIP1 and tracrRNA-NTC solutions. After 20-minute incubation, the mixtures were added to appropriate wells for a final concentration of 2 uL/mL Dharmafect, 25 nM tracrRNA, and 25 nM VCPIP1 (6.25 nM of each guide) or 25 nM NTC. Cells were incubated overnight at 37 °C. Media was aspirated and replaced after 18 hours. Cells were permitted to grow to confluency for 2 days, upsized to 6-well plates (Corning, 3516). A sample of cells were harvested, lysed, and resolved by SDS-PAGE and analyzed by Western Blot as above to assess VCPIP1 knock-out (Bethyl, A302-933A-M) with GAPDH control (Cell Signaling Technology, 97166). Monoclonal VCPIP1 KO cells were obtained through a limiting dilution plating 0.5 cells/well in a 96-well plate (Corning, 3598), with monoclonal populations being upsized to 24-well plates (Corning, 3524), with samples being taken when confluent to identify complete VCPIP1 knock out populations.

#### Golgi staining and imaging

A2058 cells, WT and KO, were grown to 70-80% confluency and then split (DMEM + 10% FBS) in 6-well plates (Falcon, 353046) to 0.4×10^6^ density in 2 mL per well. Cells were transfected the next day with a mixture of 150 µL of Opti-MEM (Gibco, 31985-070), 9 µL FuGENE ® HD Transfection Reagent (Promega, E2311) and 3 ug of DNA in a drop-by-drop manner. Media was changed the next day, and the cells were incubated for 48 hours post transfection. The cells were split into two samples post harvesting – one for confirming transfection efficiency by western blot analysis and one for qualitative analysis using confocal microscopy.

For western blot analysis, 10^6^ cells were washed in PBS, lysed in RIPA Lysis and extraction buffer (G Biosciences, 768-489) with cOmplete Mini protease inhibitor cocktail tablets (Roche, 11836153001), and the supernatant were transferred onto PVDF membranes using an iBlot 2 dry blotting system (Thermo Fisher Scientific, IB21001). Rabbit anti-VCPIP1 primary antibody (BETHYL, A302-933A) was used to detect the VCPIP1 in WT and KO conditions and the blots were imaged using LI-COR Odessey CLx detecting a donkey anti-rabbit secondary antibody (LI-COR, 926-32213). The blot demonstrated that the transfection of all the constructs was successful within the A2058 KO cell line **(Extended Data Fig. 6a**).

For qualitative analysis, 100 µL of harvested cells from each condition were cytospun (400 rpm, 5 minutes) onto slides for staining (Enzo GOLGI-ID ® Green Assay Kit, ENZ-51028-K100). Cells were protected from light to prevent photobleaching. A hydrophobic barrier pen (invitrogen, R3777) was used to create a lipid barrier around the cells. The cells were initially washed with 1X Assay buffer (GOLGI-ID kit) and then fixed with 4% of 16% paraformaldehyde solution (Electron Microscopy Sciences, 15710) at room temperature for 10 minutes before staining the cells for detection of Golgi bodies using the manufacture’s manual for “Staining Adherent Fixed Cells”. The stained cells were mounted with DPX Mountant for histology (Sigma-Aldrich, 06522-100mL) using #1.5 coverslips (Scientific Laboratory Supplies LTD, MIC3246) and left to dry in the dark for 30 minutes at room temperature and stored at 4°C until imaging.

The mounted slides were analyzed (ZEN Desktop software, (RRID:SCR_013672)) using a Zeiss 980 confocal microscope with an Airyscan 2 detector enabling super-resolution and high-speed imaging^57^. We collected Z-stacks with Airyscan of all the conditions to eliminate Z-height bias. Using Airyscan, Z-stacks (0.15 µm interval) were taken of all the conditions using the 63X 1.4 oil objective. Airyscan processed images were imported into Fiji^58^ (ImageJ) and further processed using Z-stack max intensity projection to observe areas where fragmentation was evident. Channels from the final image were split to see both DAPI and GOLGI-ID stain independently and then merged. All images were imaged and processed with the same parameters.

