## Extended Data Figures for "Ternary complex formation of VCP, VCPIP1 and p47 facilitates Golgi reassembly"

### Extended Data Figure 1: Stuctural characterization of VCP-VCPIP1 complex.

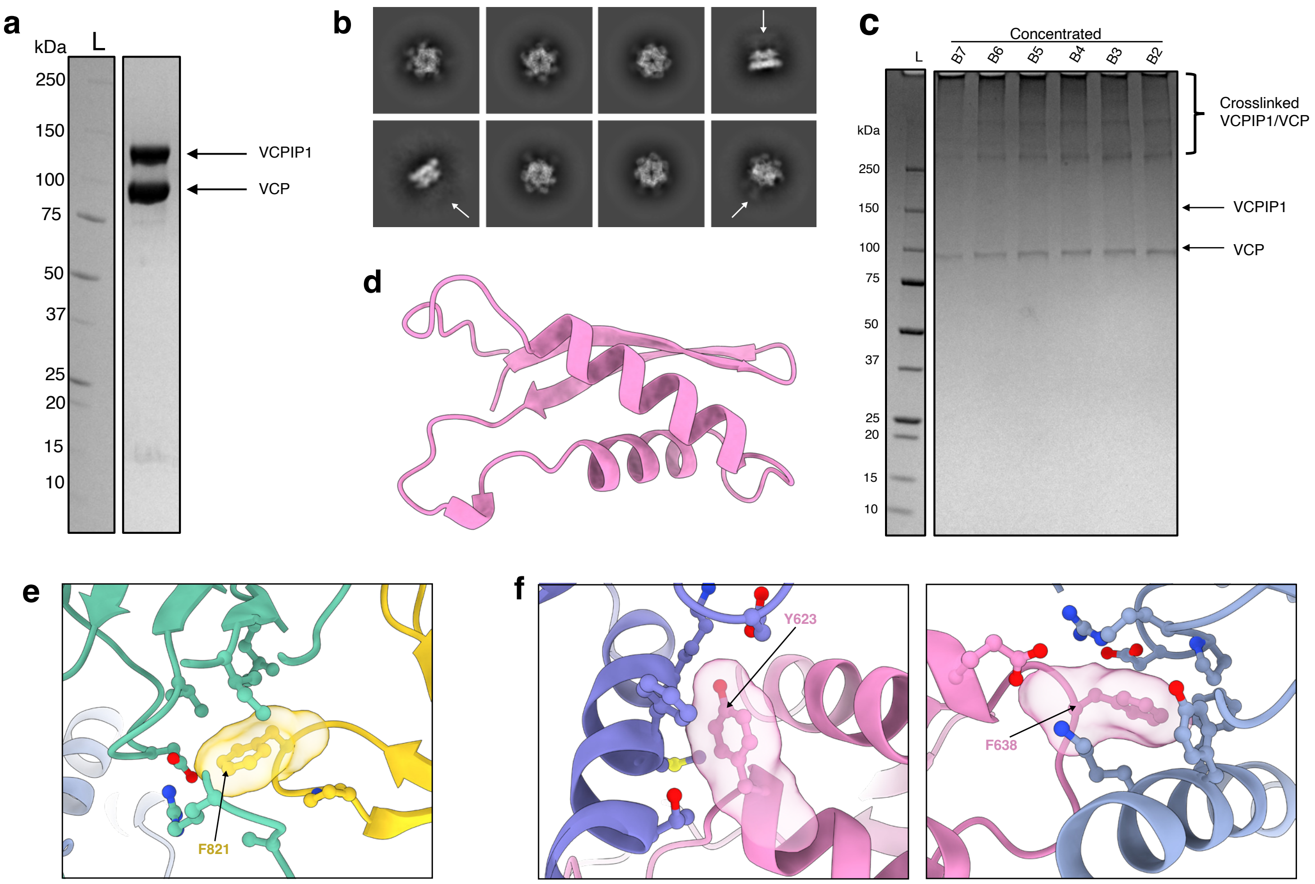

### Extended Data Figure 2. Cryo-EM processing workflow for the VCP-VCPIP1 complex.

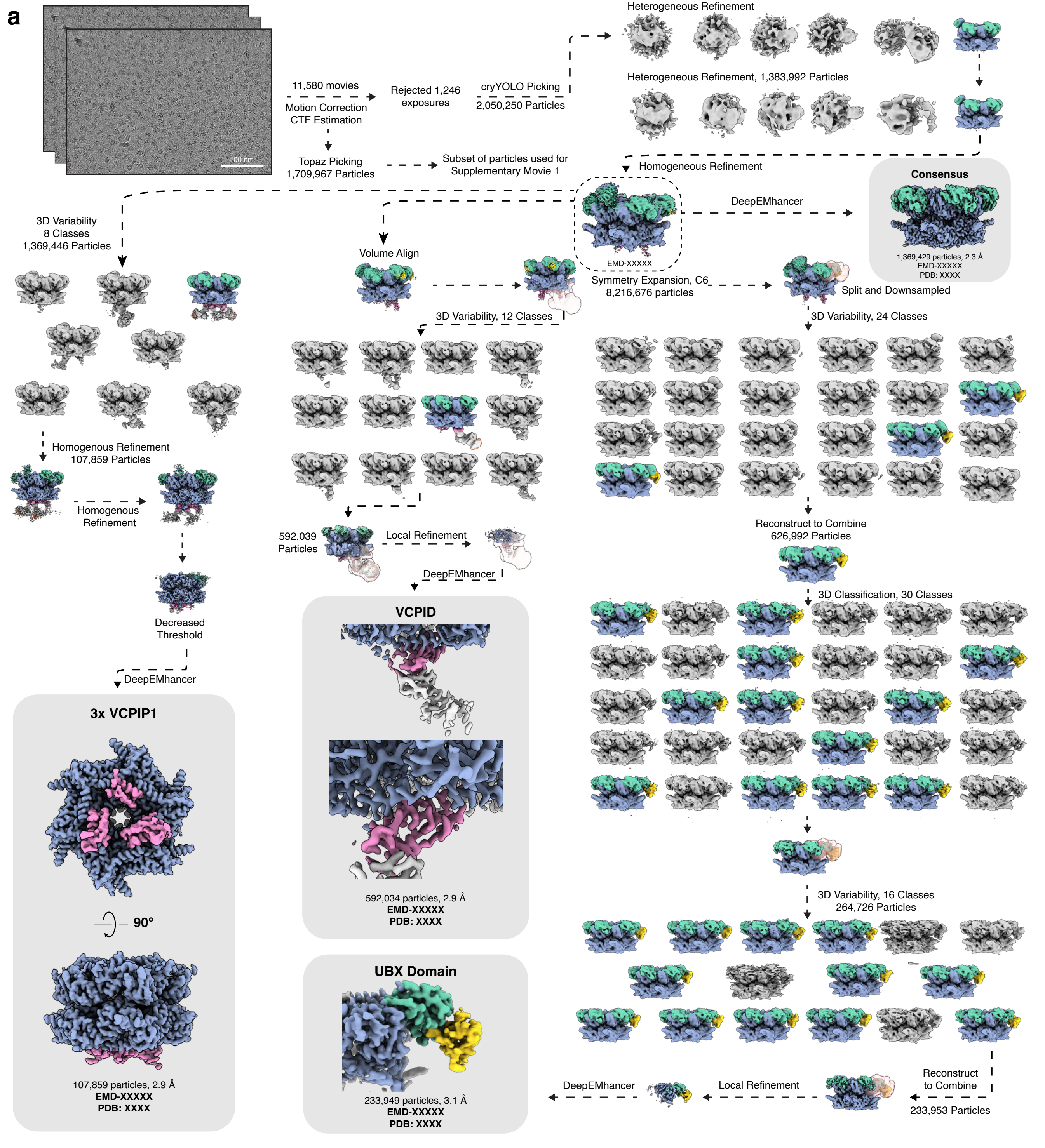

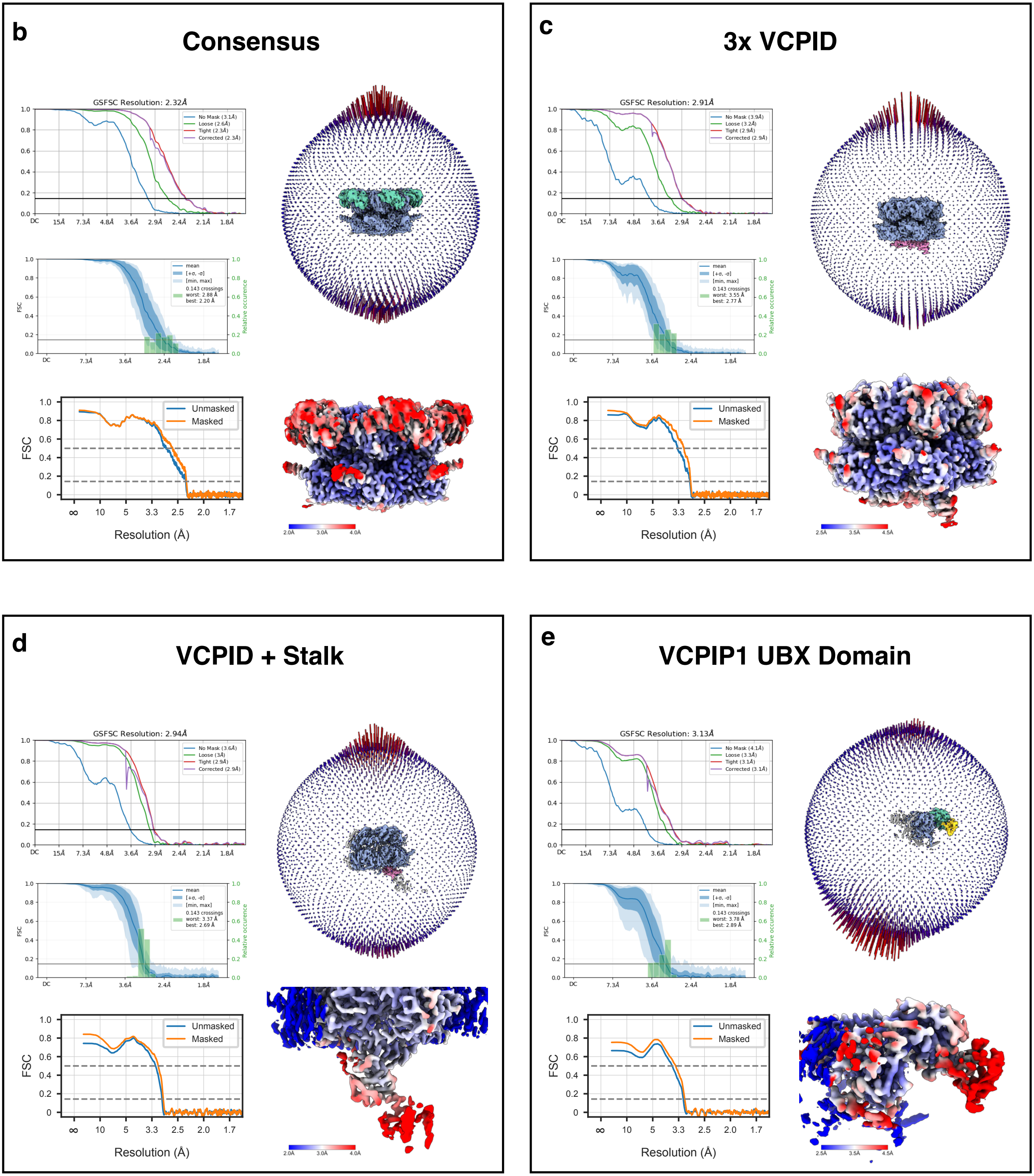

b
Consensus
GSFSC Resolution: 2.32Å
No Mask (3.1Å)
Loose (2.6Å)
Tight (2.3Å)
Corrected (2.3Å)
mean
[+σ, -σ]
[min, max]
0.143 crossings
worst: 2.88 Å
best: 2.20 Å
Relative occurence
FSC
Unmasked
Masked
Resolution (Å)
2.0Å
3.0Å
4.0Å
c
3x VCPID
GSFSC Resolution: 2.91Å
No Mask (3.9Å)
Loose (3.2Å)
Tight (2.9Å)
Corrected (2.9Å)
mean
[+σ, -σ]
[min, max]
0.143 crossings
worst: 3.55 Å
best: 2.77 Å
Relative occurence
FSC
Unmasked
Masked
Resolution (Å)
2.5Å
3.5Å
4.5Å
d
VCPID + Stalk
GSFSC Resolution: 2.94Å
No Mask (3.6Å)
Loose (3Å)
Tight (2.9Å)
Corrected (2.9Å)
mean
[+σ, -σ]
[min, max]
0.143 crossings
worst: 3.37 Å
best: 2.69 Å
Relative occurence
FSC
Unmasked
Masked
Resolution (Å)
2.0Å
3.0Å
4.0Å
e
VCPIP1 UBX Domain
GSFSC Resolution: 3.13Å
No Mask (4.1Å)
Loose (3.3Å)
Tight (3.1Å)
Corrected (3.1Å)
mean
[+σ, -σ]
[min, max]
0.143 crossings
worst: 3.78 Å
best: 2.89 Å
Relative occurence
FSC
Unmasked
Masked
Resolution (Å)
2.5Å
3.5Å
4.5Å

### Extended Data Figure 3: DUB activity assay using ubiquitin rhodamine 110 (Ub-Rho) as substrate.

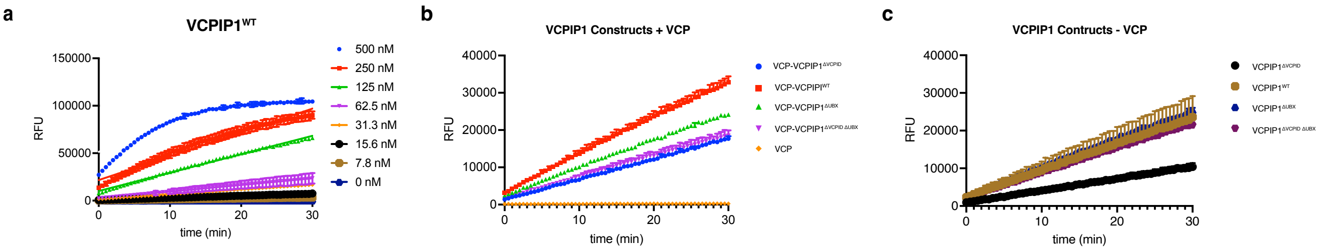

### Extended Data Figure 4: Structural characterization of VCP-VCPIP1-p47 complex.

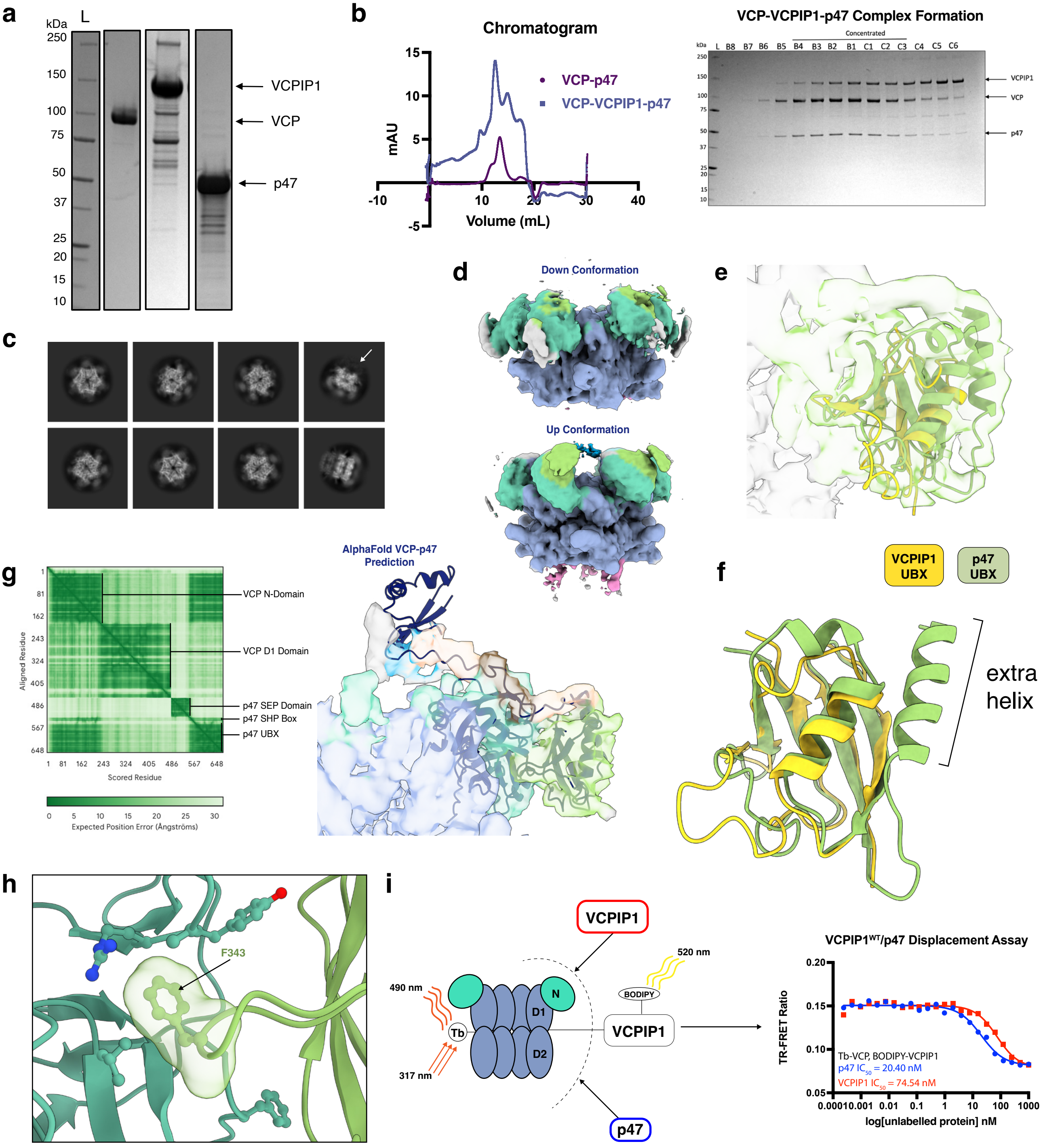

### Extended Data Figure 5. Cryo-EM processing workflow for the VCP-VCPIP1-p47 complex.

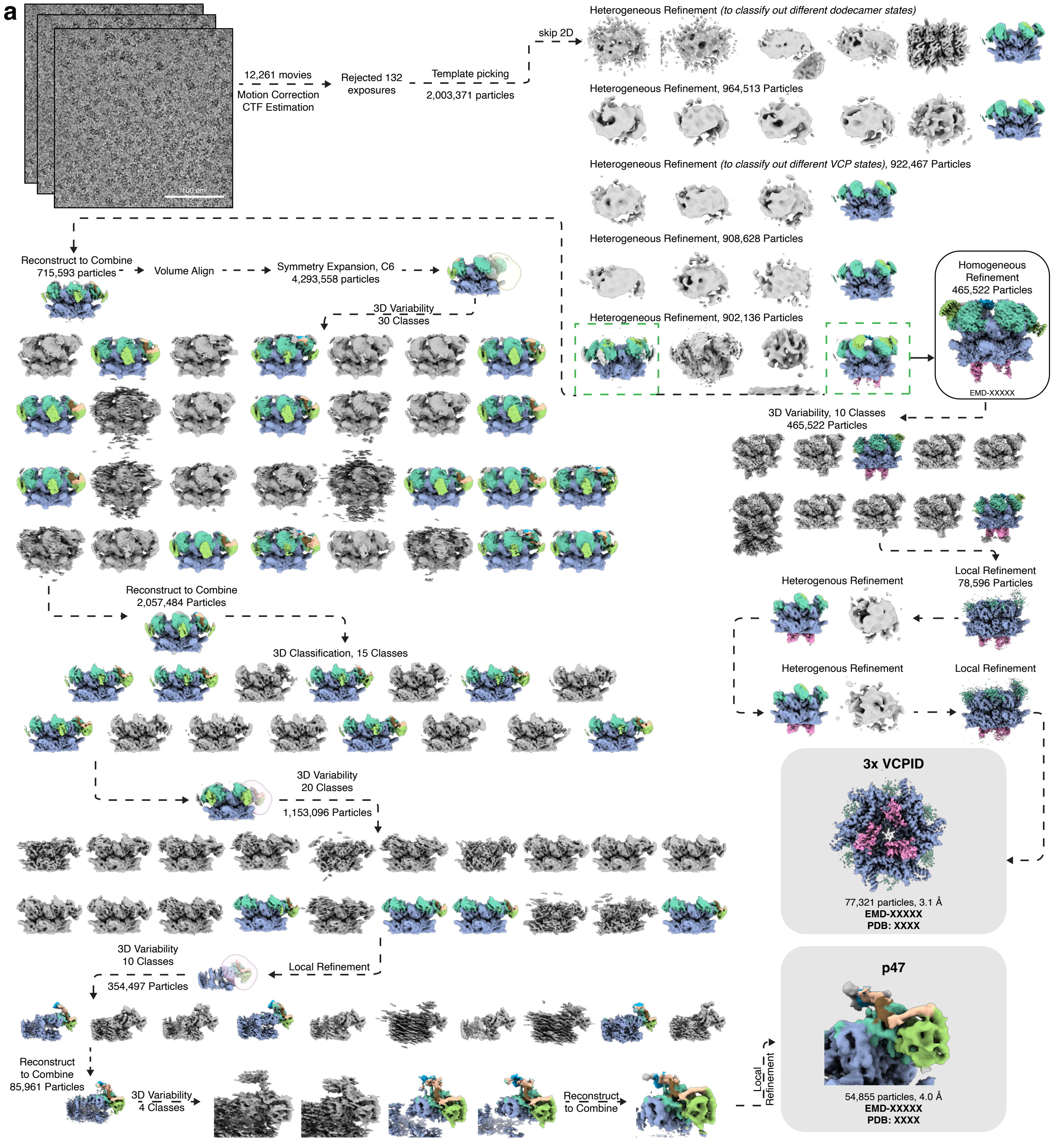

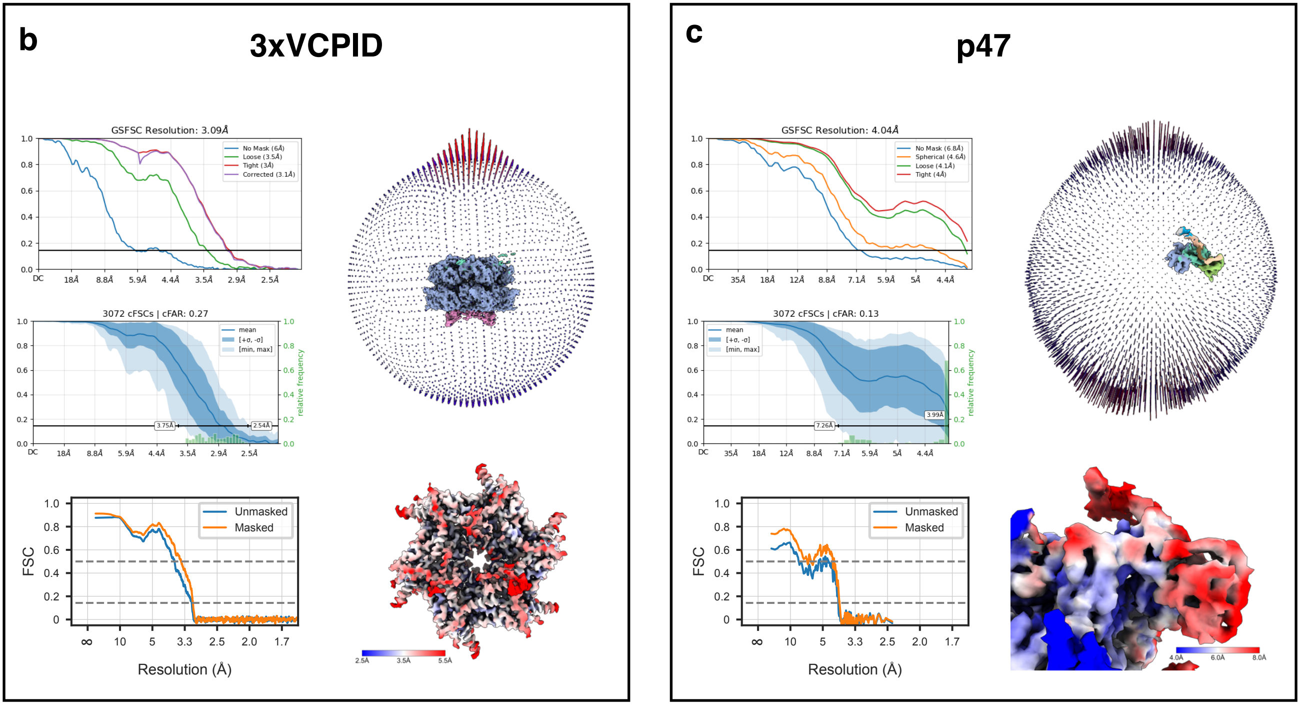
b
3xVCPID
GSFSC Resolution: 3.09Å
No Mask (6Å)
Loose (3.5Å)
Tight (3Å)
Corrected (3.1Å)
3072 cFSCs | cFAR: 0.27
mean
[+σ, -σ]
[min, max]
relative frequency
3.75Å
2.54Å
Unmasked
Masked
FSC
Resolution (Å)
2.5Å
3.5Å
5.5Å
c
p47
GSFSC Resolution: 4.04Å
No Mask (6.8Å)
Spherical (4.6Å)
Loose (4.1Å)
Tight (4Å)
3072 cFSCs | cFAR: 0.13
7.26Å
3.99Å
4.0Å
6.0Å
8.0Å

### Extended Data Figure 6: Analysis of Golgi reassembly with VCPIP1 domain truncations.

**a**

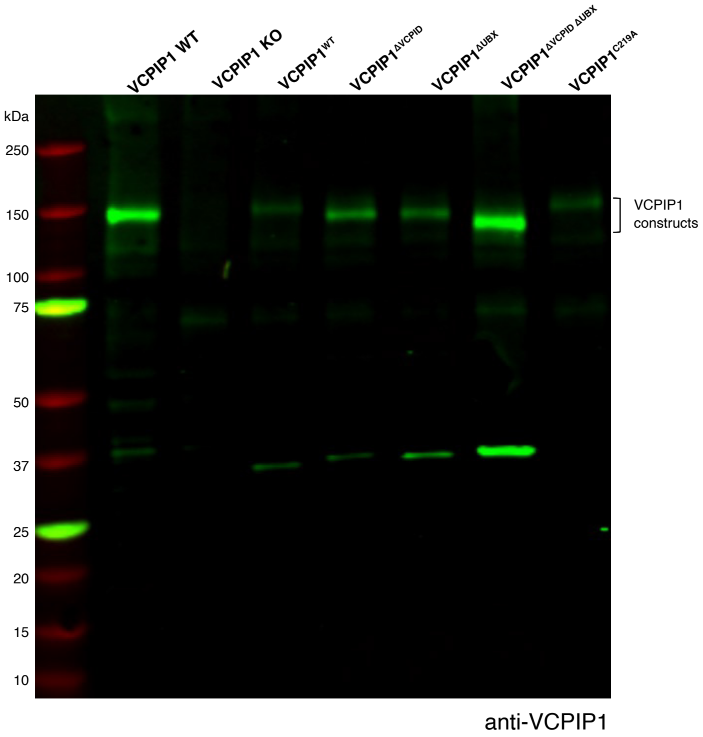

### Extended Data Figure 7. Uncropped SDS-PAGE and western blots.

Extended Data Fig. 1b

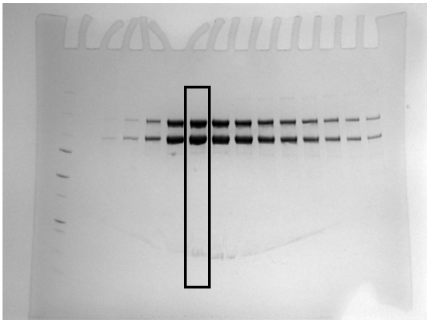

Extended Data Fig. 1c

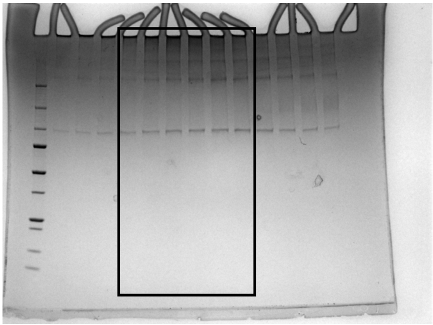

Extended Data Fig. 4a (left)

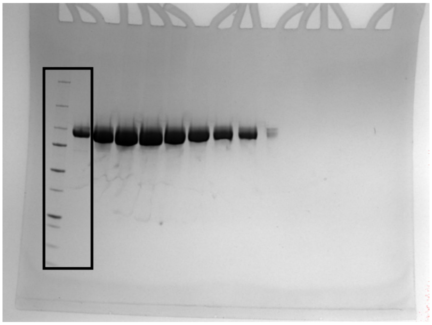

Extended Data Fig. 4a (middle)

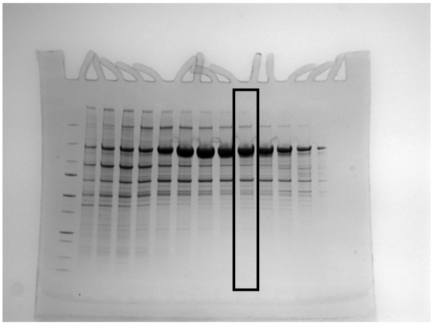

Extended Data Fig. 4a (right)

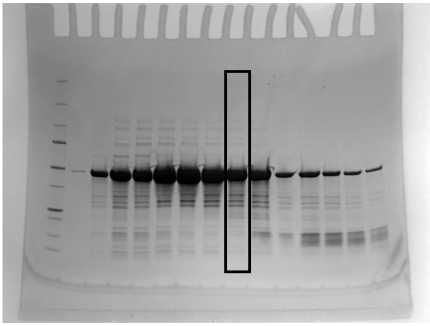

Extended Data Fig. 4b

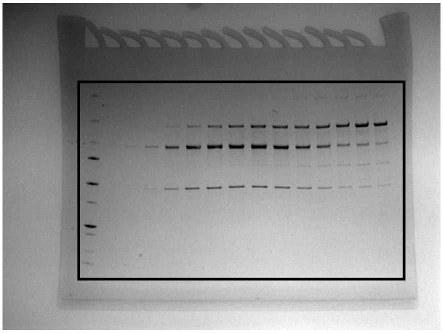

Extended Data Fig. 6

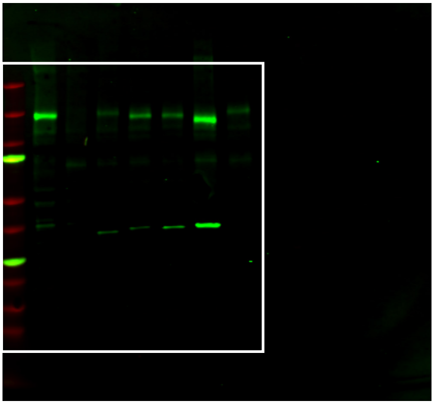
