## Supplementary material for "Ternary complex formation of VCP, VCPIP1 and p47 facilitates Golgi reassembly": Tables

### Table 1: Cryo-EM data collection, refinement and validation statistics of VCP-VCPIP1 Complex

| Data Set 1 – VCP-VCPIP1 Complex | | | | | |
|---|---|---|---|---|---|
| Microscope | Titan Krios (Thermo Fischer Scientific) | | | | |
| Voltage (kV) | 300 | | | | |
| Camera | Gatan K3 | | | | |
| Magnification | 105,000 | | | | |
| Pixel size (Å) | 0.83 | | | | |
| Total electron exposure (e–/ Å$^2$) | 53.69 | | | | |
| Number of frames (no.) | 50 | | | | |
| Defocus range (µm) | -0.9 - -2.2 | | | | |
| Data Collection Software | SerialEM (v. 3.8.6) | | | | |
| Energy filter slit width (eV) | 20 | | | | |
| Micrographs collected (no.) | 11,580 | | | | |
| Micrographs used (no.) | 10,334 | | | | |
| Total extracted particles (no.) | 2,050,250 | | | | |
| | - | Map 1 | Map 2 | Map 3 | Map 4 |
| | - | Consensus | 3xVCPID | VCPID | UBX |
| EMDB accession code | EMD-XXXXX | EMD-XXXXX | EMD-XXXXX | EMD-XXXXX | EMD-XXXXX |
| PDB accession code | - | XXXX | XXXX | XXXX | XXXX |
| Final particles used (no.) | 1,369,446 | 1,369,429 | 107,859 | 592,034 | 233,949 |
| Symmetry | C1 | C1 | C1 | C1 | C1 |
| Map resolution (Å, FSC 0.143) | 2.9 | 2.3 | 2.9 | 2.9 | 3.1 |
| Resolution range (Å) | 2.5 – 6.4 | 2.0 - 5.5 | 1.8 - 7.9 | 2.5 – 6.5 | 2.7 – 6.4 |
| Refinement Package | phenix.real_space_refine | | | | |
| **Model composition**<br>**Non-hydrogen atoms (no.)**<br>**Protein residues (no.)** | - | <br>33,395<br>4,257 | <br>27,895<br>3,563 | <br>6,872<br>872 | <br>4,117<br>523 |
| Model-to-Map CC | | 0.84 | 0.85 | 0.81 | 0.81 |
| Model-to-Map, FSC (Å, FSC 0.5) | | | | | |
| *B* factors (Å$^2$)<br>Protein (mean)<br>Water | | <br>86.22<br>- | <br>86.29<br>- | <br>87.83<br>- | <br>97.96<br>- |
| **R.m.s. deviations**<br>**Bond lengths (Å)**<br>**Bond angles (°)** | - | <br>0.002<br>0.460 | <br>0.003<br>0.476 | <br>0.003<br>0.502 | <br>0.003<br>0.510 |
| **Validation**<br>**MolProbity score**<br>**Clashscore**<br>**Rotamer Outliers (%)**<br>**CaBLAM outliers (%)** | - | <br>1.61<br>6.78<br>1.49<br>2.21 | <br>1.81<br>7.03<br>1.98<br>1.80 | <br>1.93<br>8.60<br>1.91<br>2.88 | <br>1.92<br>10.59<br>1.76<br>3.75 |
| **Ramachandran plot (%)**<br>**Favored**<br>**Allowed**<br>**Disallowed** | - | <br>97.48<br>2.49<br>0.02 | <br>96.80<br>3.17<br>0.03 | <br>96.24<br>3.76<br>0.00 | <br>96.89<br>2.91<br>0.19 |

#### Table 2: Cryo-EM data collection, refinement and validation statistics of VCP-VCPIP1-p47 Complex

| Data Set 2 – VCP-VCPIP1-p47 Complex | | | |
|---|---|---|---|
| Microscope | Titan Krios (Thermo Fischer Scientific) | | |
| Voltage (kV) | 300 | | |
| Camera | Falcon4 | | |
| Magnification | 165,000 | | |
| Pixel size (Å) | 0.736 | | |
| Total electron exposure (e–/ Å$^2$) | 49.22 | | |
| Number of frames (no.) | 49 | | |
| Defocus range (µm) | -0.8 - -2.2 | | |
| Data Collection Software | EPU (v 3.7) | | |
| Energy filter slit width (eV) | 10 | | |
| Micrographs collected (no.) | 12,261 | | |
| Micrographs used (no.) | 12,129 | | |
| Total extracted particles (no.) | 2,003,371 | | |
| | | Map 1 | Map 2 |
| | | 3xVCPID | p47 UBX |
| EMDB accession code | EMD-XXXXX | EMD-XXXXX | EMD-XXXXX |
| PDB accession code | - | XXXX | XXXX |
| Final particles used (no.) | 465,522 | 77,321 | 54,855 |
| Symmetry | C1 | C1 | C1 |
| Map resolution (Å, FSC 0.143) | 4.0 | 3.1 | 4.0 |
| Resolution range (Å) | 4.5 – 7.6 | 2.8 – 9.1 | 4.5 - 9.8 |
| Refinement Package | phenix.real_space_refine | | |
| **Model composition**<br>**Non-hydrogen atoms (no.)**<br>**Protein residues (no.)** | - | 28,483<br>3,641 | 4,314<br>554 |
| Model-to-Map CC | | 0.80 | 0.74 |
| Model-to-Map, FSC (Å, FSC 0.5) | | | |
| *B* factors (Å$^2$)<br>Protein (mean)<br>Water | - | 90.93<br>- | 145.09 |
| **R.m.s. deviations**<br>**Bond lengths (Å)**<br>**Bond angles (°)** | - | 0.002<br>0.456 | 0.003<br>0.730 |
| **Validation**<br>**MolProbity score**<br>**Clashscore**<br>**Rotamer Outliers (%)**<br>**CaBLAM outliers (%)** | - | 1.81<br>6.66<br>2.14<br>2.68 | 1.83<br>11.85<br>0.00<br>4.09 |
| **Ramachandran plot (%)**<br>**Favored**<br>**Allowed**<br>**Disallowed** | - | 96.83<br>3.17<br>0.00 | 96.34<br>3.66<br>0.00 |
